# Induction of Meiosis from Human Pluripotent Stem Cells

**DOI:** 10.1101/2024.05.31.596483

**Authors:** Merrick Pierson Smela, Jessica Adams, Carl Ma, Laura Breimann, Ursula Widocki, Toshi Shioda, George M. Church

## Abstract

An *in vitro* model of human meiosis would accelerate research into this important reproductive process and development of therapies for infertility. We have developed a method to induce meiosis starting from male or female human pluripotent stem cells. We demonstrate that DNMT1 inhibition, retinoid signaling activation, and overexpression of regulatory factors (anti-apoptotic BCL2, and pro-meiotic HOXB5, BOLL, or MEIOC) rapidly activates meiosis, with leptonema beginning at 6 days, zygonema at 9 days, and pachynema at 12 days. Immunofluorescence microscopy shows key aspects of meiosis, including chromosome synapsis and sex body formation. The meiotic cells express genes similar to meiotic oogonia *in vivo*, including all synaptonemal complex components and machinery for meiotic recombination. These findings establish an accessible system for inducing human meiosis *in vitro*.

## Main Text

All sexually reproducing species rely on meiosis to produce haploid gametes from diploid germ cells. To date, the most detailed studies of meiosis have taken place in non-human organisms, due to the lack of a reliable *in vitro* model of human meiosis, as well as technical and ethical barriers to obtaining meiotic cells from humans. Therefore, a method of inducing meiosis in cultured human cells could greatly advance the study of this crucial reproductive process, and could also lead to new therapies for people with infertility.

Research on animals such as mice has revealed important characteristics of mammalian meiosis, including requirements of erasing DNA methylation (*1*), as well as retinoic acid and BMP signaling from gonadal somatic cells (*2*). Recent studies have demonstrated the initiation of meiosis in mouse cells *in vitro* (*2–8*), even producing viable offspring from the resulting gametes (*3*, *4*). However, studies attempting to initiate meiosis in human cells have been less successful. These studies based their main conclusions on the production of haploid (1N 1C) cells as assessed by flow cytometry for DNA content (*9–12*). However, this approach has two important flaws (*13*). First, the 1N 1C state is non-physiological in eggs since meiosis is not completed until after fertilization. Second, dead and dying cells with fragmented nuclei can have reduced DNA content, leading to false positives in this assay. Some studies also examined the expression of the meiotic markers SYCP3 and γH2AX (*11*, *12*, *14*, *15*), but did not convincingly show the expected localization of these proteins during the stages of meiosis, and our attempts to reproduce these protocols were unsuccessful (Table S1).

Here, we present an *in vitro* model of meiosis from human induced pluripotent stem cells (hiPSCs). By screening conditions for activating the expression of meiotic genes, we found that DNMT1 inhibition, retinoic acid receptor activation, and overexpression of anti-apoptosis and pro-meiosis factors can rapidly initiate meiosis in male and female hiPSCs. We show that this method generates cells corresponding to the leptotene, zygotene, and pachytene stages of meiosis, and that these cells have gene expression similar to meiotic germ cells *in vivo*. Overall, our method will be a useful tool for researchers studying human meiosis, and with further optimization may allow the production of human gametes *in vitro*.

### Barcode enrichment screening of candidate meiosis-promoting factors

We began by analyzing previously published scRNAseq data of human fetal gonads, which contain a variety of cell types (*16*), including pre-meiotic STRA8+ oogonia and fully meiotic oogonia/oocytes (Fig. 1A). We confirmed *REC8* as a reliable marker for early meiotic cells, and *SYCP3* for late meiotic cells. Using previously constructed male and female DDX4-tdTomato reporter hiPSCs (*17*), we engineered dual reporter lines for DDX4-tdTomato/REC8-mGreenLantern (D4TR8G) and DDX4-tdTomato/SYCP3-mGreenLantern (D4TS3G) (Fig. S1). We validated these lines by whole genome sequencing (*18*), and by CRISPRa and flow cytometry.

**Fig. 1.**
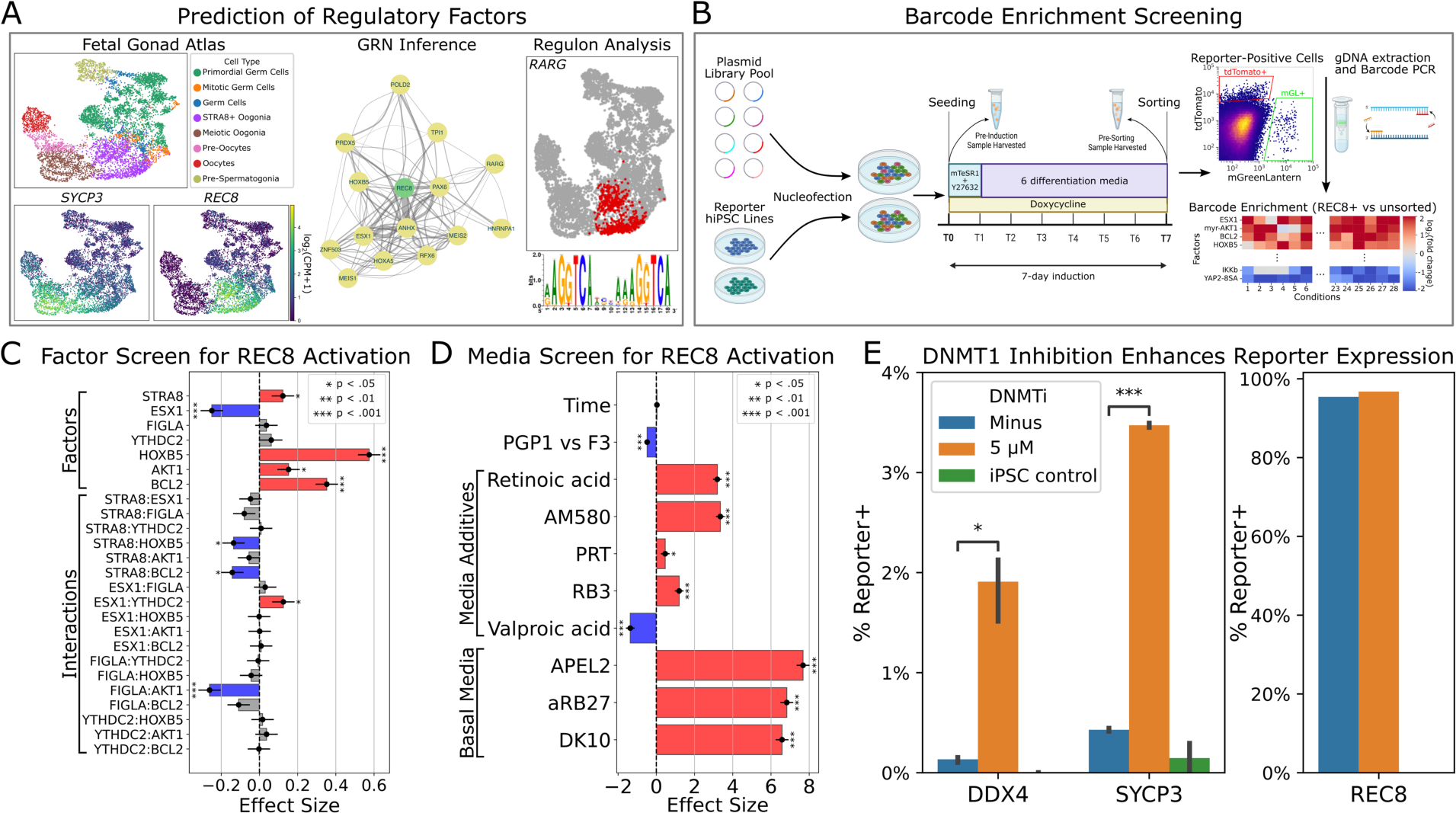
Screening of regulatory factors for reporter activation. (A) Prediction of regulatory factors based on fetal gonad scRNAseq data. (B) Barcode enrichment screening in reporter iPSCs (see Methods). (C) Fractional factorial screen (32 combinations, each tested in 2 cell lines) of seven top factors for REC8 activation. Effect sizes and significance were calculated using a linear model on logit-transformed data. (D) Screen of media and additives for REC8 activation (see Methods). Effect sizes and significance were calculated as above. (E) Effects of DNMT1i treatment on expression of DDX4, SYCP3, and REC8 reporters (n=4).

Next, we chose 78 candidate meiosis-promoting factors based on scRNAseq analysis and previous literature (see Methods). We cloned these into a barcoded PiggyBac transposon plasmid for doxycycline-inducible expression. We integrated the library into the reporter hiPSCs, activated expression, and sorted reporter-positive cells after seven days of induction (Fig. 1B). We tested low-copy and high-copy integration, as well as a variety of different culture media (see Methods). Comparing barcode frequencies between reporter-positive and unsorted populations, we found several factors consistently enriched in REC8+ cells (Fig. S2), although results for the other reporters were noisy due to low cell yield.

### Optimization of REC8 activation

Based on the barcode enrichment results, we narrowed down our library to sixteen factors and tested these individually for activation of REC8 and SYCP3 expression (Fig. S3). BCL2, HOXB5, and myr-AKT1 all slightly activated REC8, although no factors activated SYCP3. We next performed a combinatorial screen of seven factors, and found that BCL2, HOXB5, STRA8, and myr-AKT1 all significantly promoted REC8 expression (Fig. 1C). Using these top four factors, we tested different media compositions, supplements, and induction times (Fig. 1D). The best-performing medium was APEL2, and retinoids (retinoic acid and AM580) significantly increased REC8 expression. The PRC1 inhibitors RB3 and PRT4164 caused a small increase in REC8 expression, but this was associated with extensive toxicity. Valproic acid was also toxic, and significantly decreased REC8 expression. There was no significant change in REC8 expression between 6, 7, and 8 days of induction.

### DNMT1 inhibition upregulates meiotic markers

Using the top factors and differentiation medium, we could induce REC8+ cells at nearly 100% efficiency, but cells still lacked SYCP3 expression. We reasoned that overexpressing meiosis-promoting factors might not be sufficient, and that downregulation of meiosis-inhibiting factors might be required. Therefore, we tested CRISPRi knockdown of ten epigenetic factors. Knockdown of *DNMT1* resulted in a small upregulation of SYCP3 (Fig. S4). In previous work, we used a noncovalent DNMT1 inhibitor, GSK3484862, to erase DNA methylation and establish an oogonia-like epigenetic state (*17*). Treatment with this inhibitor resulted in a significant increase in expression of SYCP3, as well as DDX4 (Fig. 1E).

### scRNAseq screening identifies meiotic cells and associated factors

With DNMT1 inhibition, retinoid treatment, and overexpression of BCL2, HOXB5, and STRA8, we observed activation of REC8, SYCP3, and DDX4. However, we wanted to take a broader view of the gene expression in our cells, and see if expressing any additional factors could drive the cells closer to meiosis. Therefore, we generated iPSC populations containing integrated expression vectors for BCL2, HOXB5, and STRA8 under hygromycin selection, as well as a pool of 88 other candidate regulatory factors under puromycin selection (Supplementary Table 2). Following our induction protocol, we performed scRNAseq on sorted reporter-positive cells as well as unsorted cells (Fig. 2A, and see Methods).

**Fig. 2.**
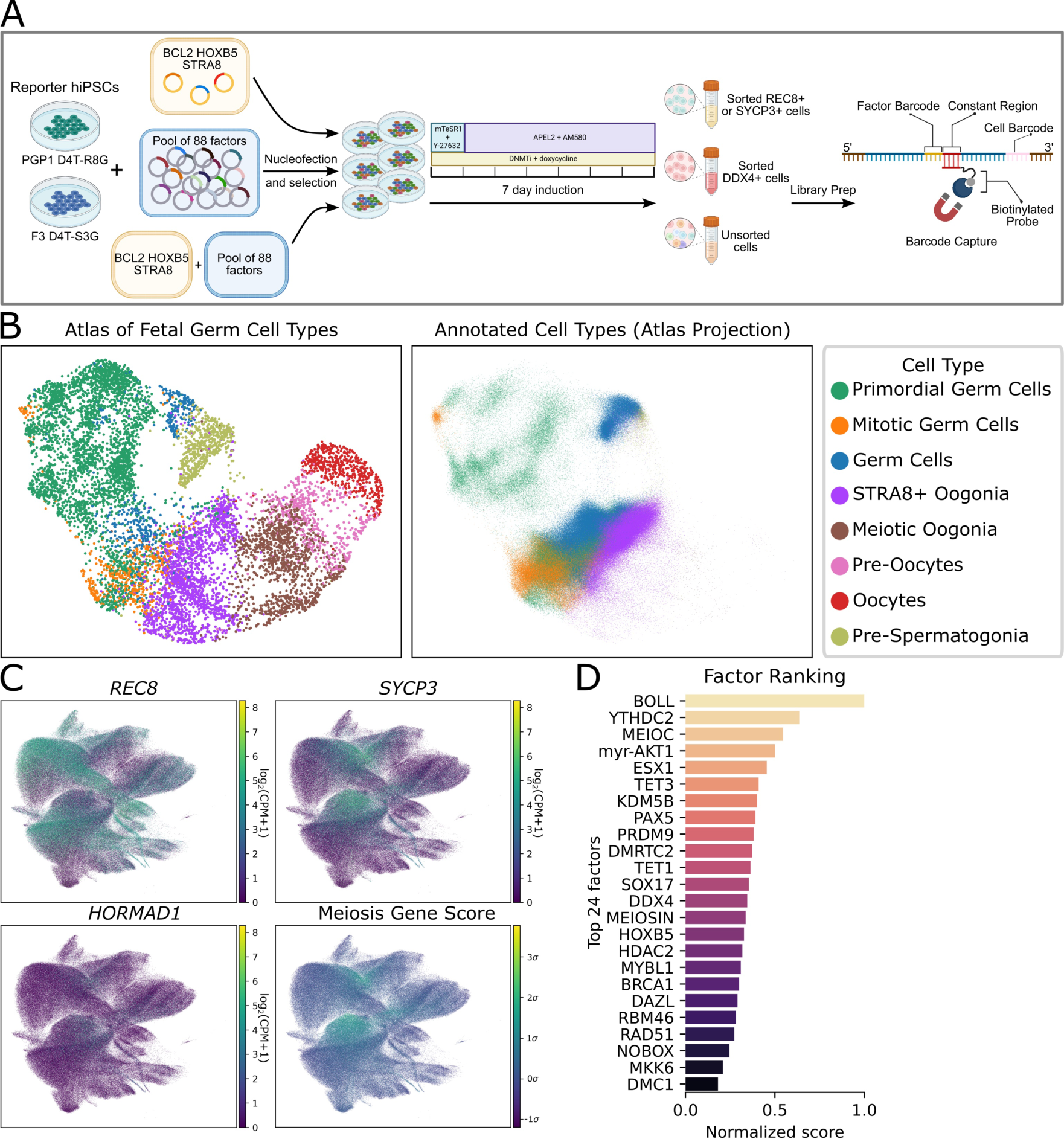
scRNAseq identifies factors promoting meiotic cell identity. (A) scRNAseq differentiation, library prep, and barcode capture (see Methods). (B) Cell type annotation based on the fetal germ cell atlas. (C) Expression of selected meiosis marker genes, as well as the meiosis gene score calculated from expression of 19 meiosis-specific genes. (D) Factor ranking based on barcode overrepresentation in cells with meiotic gene expression.

We first investigated whether any meiotic cells were present in our samples. Leveraging the fetal gonad scRNAseq dataset (*16*), we performed cell type annotation (Fig. 2B). The majority of our cells were classified as pre-meiotic oogonia, and small fraction of cells were annotated as fully meiotic. The proportion of these cells was greatest (0.8%) in sorted SYCP3+ samples. As another way of looking for meiotic cells, we constructed a score based on expression of meiosis-specific genes. In addition to *REC8* and *SYCP3*, the markers for which we sorted, we also observed expression of other essential meiosis genes, such as *HORMAD1*, in a smaller fraction of cells (Fig. 2C). We chose a list of nineteen meiosis-specific genes (Supplementary Table 3) and scored cells based on their expression. Out of 646,493 cells in our dataset, 1,276 cells had a gene score >4σ, compared with an expected 20 cells assuming random gene expression.

Next, we asked which regulatory factors were responsible for generating these meiotic cells. We performed a hybridization-based capture to enrich our scRNAseq library for barcode sequences and identified at least one expressed barcode in 91% of our cells. We then examined which factors were overrepresented in meiotic cells (defined using cell type annotation or gene scoring) versus pre-meiotic cells. We chose the top 24 for subsequent screening (Fig. 2D).

### Optimization of factors for inducing meiosis

We next expressed each of these 24 factors along with the previous top three (BCL2, HOXB5, and STRA8). After seven days of induction, we monitored reporter activation and performed immuno-staining for the meiosis markers HORMAD1 and SYCP3. We identified four factors, BOLL, MEIOC, MEIOSIN, and myr-AKT1, as the most promising (Fig. S5A). We combined these four with the previous top three and repeated the experiment, this time analyzing a series of time points (7, 9, 13, and 16 days). Excitingly, at days 9 and 13 post-induction, we observed a few HORMAD1+ SYCP3+ cells with zygotene-like filamentous staining (Fig. S5B).

In order to narrow down which factors were responsible for inducing meiosis, we performed two rounds of combinatorial screening. In the first round, we tested 16 combinations of the initial set of seven factors (Fig. S5CDE). No combination lacking BCL2 was able to induce zygotene cells as measured by HORMAD1 filament formation. BCL2/HOXB5/BOLL and BCL2/HOXB5/myr-AKT1/MEIOC were the best individual combinations. BCL2, HOXB5, and BOLL all induced a significant increase in the number of zygotene cells. Interestingly, we found that STRA8 significantly decreased the zygotene score when overexpressed.

Finally, we generated hiPSCs constitutively expressing BCL2, and inducibly expressing HOXB5, BOLL, and MEIOC, testing all eight possible combinations of these three factors. We observed that each of the factors could induce meiosis when expressed with BCL2 (Fig. 3A). BOLL was the most efficient of the three factors we tested (Fig. 3B). In the BCL2-only control, we observed no HORMAD1+ cells and only a few occasional SYCP3+ cells.

**Fig. 3.**
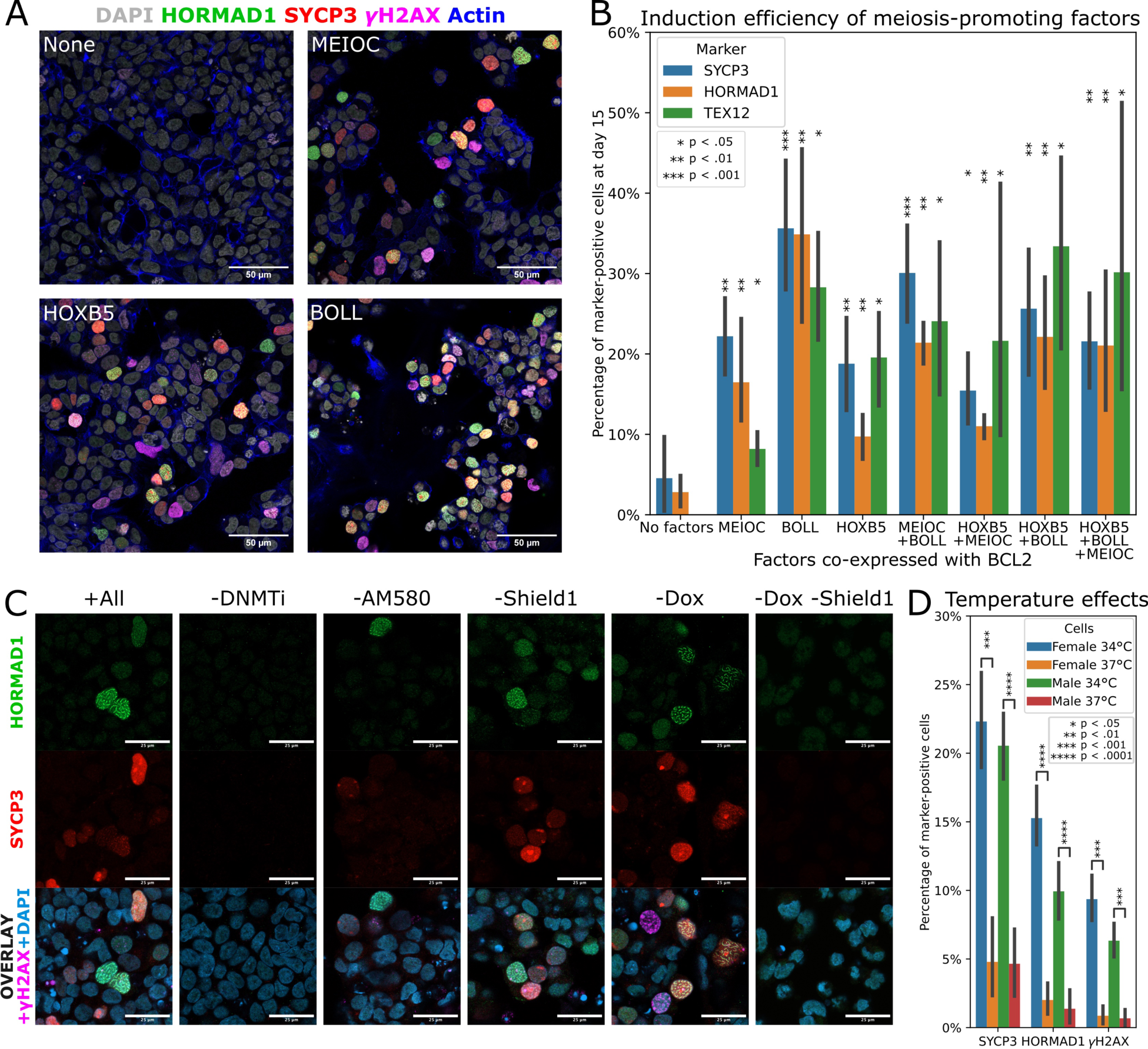
Optimization of meiosis induction. (A) Cells expressing constitutive BCL2 and Dox-inducible MEIOC, HOXB5, or BOLL were subjected to the meiosis induction protocol (see Methods) and stained for DNA (DAPI), actin (phalloidin), HORMAD1, SYCP3, and γH2AX (scale bar 50 µm). (B) From the same experiment as panel A, all eight possible combinations of the factors tested in three cell lines. Two images were analyzed per cell line and combination. (C) Cells expressing constitutive BCL2, Dox-inducible BOLL, and Shield1-inducible HOXB5 were subjected to the meiosis induction protocol omitting various factors (see Methods). Representative immunofluorescence images are shown (scale bar 25 µm). (D) Effects of performing meiosis induction in male and female hiPSCs at 34 °C or 37 °C (n = 4 samples per condition).

### DNA demethylation and retinoid stimulation are required for eGicient meiotic initiation

To further investigate the conditions necessary for meiotic initiation, we omitted different components of our protocol (Fig. 3C, Fig. S6). Without DNMT1 inhibition, meiosis was completely blocked. However, DNMT1 inhibitor could be withdrawn after the first five days without negatively affecting results. Omitting the retinoid AM580 resulted in fewer meiotic cells, but some were still present. If AM580 treatment was started later than day five, results were similarly poor. Finally, using orthogonal induction systems (Dox and Shield1) for BOLL and HOXB5 expression, we confirmed that expression of at least one of these factors was required for inducing meiosis.

### Lower temperatures enhance meiotic induction

In males, meiosis takes place in the adult testes, which are cooler than the rest of the body. Previous studies indicated that male meiosis is less efficient at 37 °C (*8*). Therefore, we tested our meiosis induction protocol at 34 °C vs. 37 °C using male and female hiPSCs. Meiosis induction was significantly enhanced at 34 °C in not only male, but also female cells (Fig. 3D, Fig. S7A). We tested 34 °C starting at either day 1 or day 3 of the induction protocol, and found that both worked equally well. Monitoring the cells for up to 21 days, we saw that cell viability declined past day 16. Interestingly, we noticed that REC8-mGreenLantern fluorescence was much weaker at 34 °C compared to 37 °C, whereas SYCP3-mGreenLantern and DAZL-mGreenLantern were equally bright (Fig. S7B).

### Identification of stages of meiosis

To analyze which stages of meiosis were present in our cells, we performed co-staining for HORMAD1, which marks the chromosome axis and is removed from synapsed chromosomes during pachynema, SYCP3, which marks the lateral elements of the synaptonemal complex, and γH2AX, which marks recombination-related DNA damage in leptonema and zygonema, and the sex body (unsynapsed XY chromosomes) of male cells in pachynema and diplonema (*19*). By day 12 of our induction protocol, three stages of meiosis (leptonema, zygonema, and pachynema) were visible. A representative image of these three stages is shown in Fig. 4A. The leptotene cell (labeled a) has diffuse HORMAD1 and SYCP3 expression and a faint γH2AX signal. The zygotene cell (labeled b) has filamentous HORMAD1 and SYCP3, and stronger γH2AX. Additionally, the chromosomes are starting to compact, as seen in the DAPI channel. The pachytene cell (labeled c) has fully compacted chromosomes, associated with SYCP3 staining. HORMAD1 staining is much weaker, and a γH2AX positive sex body (labeled with an arrow) is visible on the nuclear periphery. A 3D z-stack of this image is provided as Supplementary Video 1. Meiotic cells also expressed cytoplasmic DDX4, and nuclear foci of the recombination marker RAD51 (Fig. S8).

**Fig. 4.**
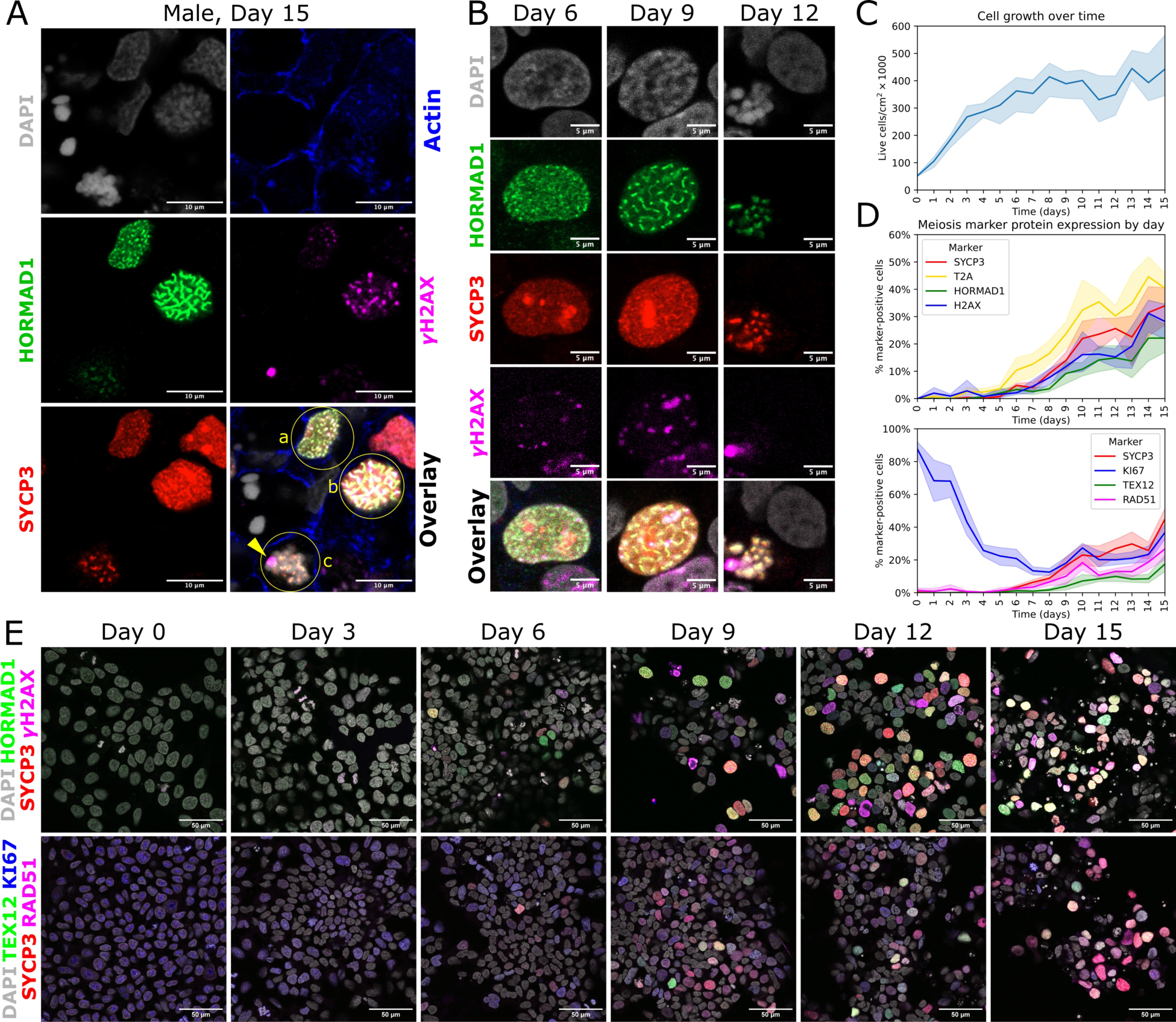
Progression of stages of meiosis over time. (A) Immuno-staining of day 15 male meiotic cells. Three stages of meiosis are visible: (a) leptonema, (b) zygonema, and (c) pachynema. A γH2AX-positive sex body, labeled with an arrow, is visible on the periphery of the pachytene nucleus. Scale bar is 10 µm. (B) Representative images of nuclei at different time points during meiosis induction in male hiPSCs. Scale bar is 5 µm. (C) Cell growth over 15 days of meiosis induction (two female and one male hiPSC line). (D) Meiosis marker protein expression over 15 days of induction (two female and one male hiPSC line, two images analyzed per line per time point). REC8 expression was measured by staining for the T2A linker peptide. (E) Representative images of meiosis marker expression over time. Scale bar is 50 µm.

### Meiotic progression over 15 days of induction

Using our optimized protocol, we investigated the progression of meiosis over time. Using three hiPSC lines (two female and one male), we measured a total of 16 timepoints per line (Fig. 4 and Supplementary Data), every 24 hours from the beginning of our induction protocol (day 0; hiPSCs) through day 15. Leptotene cells were first seen at day 6, zygotene cells were first seen at day 9, and pachytene cells were first seen at day 12 (Fig. 4B). We counted the number of live cells at each timepoint. The cells proliferated ∼8-fold over the first week of the protocol, but the number remained stable after day 8 (Fig. 4C). At the beginning of the protocol, nearly all cells were positive for KI67, a marker of proliferating cells as well as meiotic cells (*20*). As cell proliferation slowed, KI67 decreased from days 0– 7, but remained expressed in meiotic cells (Fig. 4D).

REC8-T2A was the first meiotic marker to be expressed, starting at day 6 and continuously increasing through day 11 (Fig. 4D and E). HORMAD1, SYCP3, γH2AX, and RAD51 expression followed similar trajectories, starting around days 7–8 and increasing through day 15. TEX12, which is required for full chromosome synapsis in zygonema and pachynema (*21*), was the last marker to be expressed, starting around day 9 and increasing thereafter (Fig. 4D and E). The kinetics were similar in male and female hiPSC lines.

### Gene expression dynamics during meiosis induction

We next analyzed scRNAseq data from each day of our meiosis induction protocol. Our post-filtering dataset comprised a total of 69,018 cells from one male (PGP1) and two female (F2 and F3) cell lines, and sixteen time points spanning days 0 to 15. We first performed dimensionality reduction (Fig. 5A and 5B) and examined marker gene expression (Fig. 5C, Fig. S9). Cells from the two female lines overlapped, but the male cells were largely separate (Fig. 5A). However, at later time points, the male and female lineages converged (Fig. 5B). Expression of the pluripotency marker *POU5F1* was initially high (Fig. 5C), but quickly declined and reached low levels by day 6. At intermediate time points, cells began to express gonadal germ cell markers, including *DDX4* (Fig. 5C), *DAZL, MAEL, STK31,* and *MAGE* and *PIWI* family genes (Fig. S9). Cells also expressed marker genes for meiosis, including all components of the synaptonemal complex. As observed by immunofluorescence (Fig. 4), *REC8* was one of the earliest meiosis genes expressed. By days 12-15, a subset of cells expressed late-stage pachytene recombination markers such as *MSH4* (Fig. 5C, Fig. S9). Markers for gametes (oocytes and sperm) were not highly expressed.

**Fig. 5.**
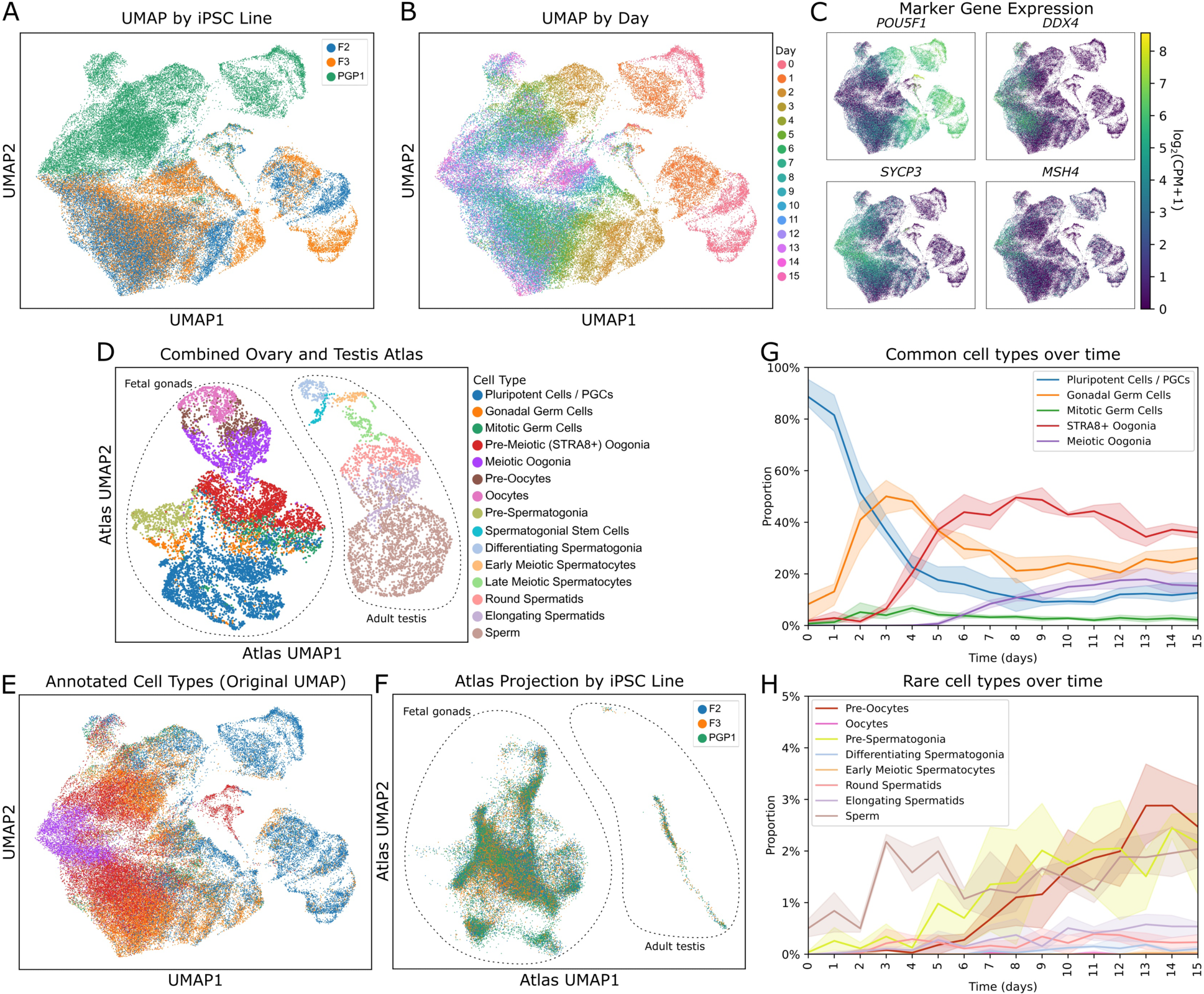
Timecourse scRNAseq analysis of meiosis induction. (A) UMAP plot, colored by hiPSC line. PGP1 is male; F2 and F3 are female. (B) UMAP plot, colored by day of sample collection. (C) UMAP plots colored by expression of marker genes for pluripotency (*POU5F1*), oogonia/gonocytes (*DDX4*), meiosis (*SYCP3*), and late-stage meiotic recombination (*MSH4*). (D) Cell types present in the combined ovary and testis reference atlas. (E) UMAP plot of annotated cell types over 15 days of meiosis induction. (F) iPSC-derived cells projected onto the atlas UMAP, colored by cell line. (G) Proportions of common (>5% abundance) cell types over time. (H) Proportions of rare (<5% abundance) cell types over time.

To compare our cells with *in vivo* germ cells, we constructed a scRNAseq atlas by combining data from human fetal gonads (containing female meiotic cells) and adult testis (containing male meiotic cells) from two previously published atlases (Fig. 5D) (*16*, *22*). We projected our cells onto the combined atlas and performed cell type annotation (Fig. 5E and 5F). This analysis showed that our cells were more similar to fetal ovarian cells, although a few cells were classified as adult testicular cells. When projected onto the atlas UMAP, cells from all three iPSC lines overlapped, and there was no clear distinction between male and female lines (Fig. 5F).

We next examined the proportions of each cell type over time, plotting common (>5% abundance) and rare (<5% abundance) cell types separately (Fig. 5G and 5H). Although the cells at early timepoints were largely annotated as primordial germ cells (PGCs), this reflects expression of marker genes such as *POU5F1* and *NANOG* which are shared between pluripotent cells and PGCs, rather than a *bona fide* PGC-like state. Indeed, our cells lacked expression of definitive PGC marker genes including *NANOS3*, *PRDM1*, *SOX17*, and *TFAP2C* (Fig. S8). Despite skipping over the PGC state, our cells transitioned through gonadal germ cell and oogonia-like states before entering meiosis (Fig. 5G). Fully meiotic cells were first present at day 6, with the proportion increasing through day 12. At later timepoints, some cells were classified as diplotene-arrested (pre-oocytes or oocytes) or post-meiotic (round spermatids, elongating spermatids, or sperm). The proportion of these cell types increased over time and reached a maximum on day 13 (Fig. 5H).

## Discussion

Here we report a reliable and rapid protocol for inducing meiosis in male and female human cells. Our method relies on overexpressing BCL2 and at least one meiosis-promoting factor. We identified HOXB5, BOLL, and MEIOC as able to perform this role. Of these, BOLL and MEIOC were previously reported as pro-meiotic (*11*, *23*). HOXB5 was known to be expressed in fetal oogonia (*16*), but its role in meiosis was not previously studied. The most likely role of BCL2 in our protocol is to prevent apoptosis resulting from DNA double strand breaks during leptonema (*24*). However, it is possible that BCL2 plays an additional role, as BCL2 alone was sufficient to upregulate REC8 (Fig. S3B). In accordance with previous studies in mice (*1*, *2*, *8*), we additionally show that DNA demethylation is required for meiotic entry, and that retinoid treatment and lower temperatures increase the efficiency.

Comparing the gene expression of our cells to *in vivo* meiotic cells, we find meiotic cells induced from both male and female hiPSCs are more similar to meiotic oogonia *in vivo*, although a small fraction (<5%) are similar to meiotic spermatocytes. Although our cells express oogonia/gonocyte markers, they do not transition through a PGC-like stage prior to meiotic entry, suggesting that this stage is not required for meiosis.

The primary limitation of our method is its low efficiency in producing pachytene and later-stage cells. We are currently using polyclonal populations of iPSCs with randomly integrated expression vectors, and switching to a system that allows precise control of transgene expression levels may improve results. Furthermore, in cultured mouse spermatogonia, meiosis has lower efficiency and fidelity compared with meiosis *in vivo* (*8*), suggesting an important role for the gonadal niche. Thus, integrating meiotic cells into recently developed ovarian organoid systems may be beneficial (*25*, *26*). Despite its modest efficiency, our current method is easily scalable and produces late-stage meiotic cells in a relatively short time (13-15 days), similar to the known duration of human meiosis (*27*).

The ability to induce meiosis using human cells *in vitro* will unlock new opportunities for science and medicine. Two examples include screening candidate male contraceptives, and using knockout hiPSCs to investigate effects of mutations. Future developments could allow the production of human gametes *in vitro*, or the generation of genetic crosses between different human cell lines. We believe that our method of inducing meiosis will greatly benefit research into this important reproductive process.

## Acknowledgements

We thank Benjamin Angulo for helpful discussions and SMAD expression plasmids used in scRNAseq screening. Portions of this research were conducted on the O2 High Performance Compute Cluster, supported by the Research Computing Group, at Harvard Medical School.

## Funding

National Institutes of Health grant F31HD108898-01A1 (MPS) Manifest Repro Grant (MPS, GC) Gift from Craig Falls (MPS, GC) Silicon Valley Community Fund grant (MPS and GC)

## Author Contributions

Conceptualization: MPS Investigation: MPS, JA, CM Resources: MPS, GMC Software: MPS, LB, UW Visualization: MPS, UW Funding Acquisition: MPS Writing–original draft: MPS Writing–review and editing: MPS, TS, GMC Supervision: TS, GMC

## Competing Interests

MPS and GMC have filed a provisional patent application on the meiosis induction protocol. A full list of GMC’s conflicts of interest can be found at https://arep.med.harvard.edu/gmc/tech.html Other authors declare that they have no competing interests.

## Data and materials availability

Sequencing data have been deposited to NCBI repositories under the accession GSE268385 [SRA accession pending]. Microscope images will be made available on Dryad following publication. Analysis code is available on Github: https://github.com/mpiersonsmela/meiosis Plasmids will be made available on Addgene following publication. All other materials, including cell lines, will be made available upon request under an MTA for noncommercial use.

## Methods

### iPSC culture

Human iPSC lines (ATCC-BXS0115, referred to as F2, ATCC-BXS0116, referred to as F3, and PGP1) were cultured in mTeSR Plus medium (Stemcell Technologies) on standard polystyrene plates coated with hESC-qualified Matrigel (Corning). hiPSCs were grown in a 37 °C 5% CO2 incubator. Passaging was performed by brief (3-5 minute) treatment with 0.5 mM EDTA and 0.25X TRYPLE Express (Gibco) in phosphate-buffered saline (PBS), followed by pipetting to break the colonies into small clumps. Cells were treated with 10 µM Y-27632 for 24 hours after each passage. Cells were tested every three months for mycoplasma using the ATCC Universal Mycoplasma Detection Kit. All cells tested negative. For experiments requiring single cell dissociation and counting, cells were harvested with Accutase and counted using trypan blue staining with a Countess II automated cell counter (Thermo Fisher).

### Generation and verification of reporter lines

Knock-in donor plasmids targeting *REC8* and *SYCP3* were constructed by Gibson assembly of 5’ and 3’ homology arms, an insert containing a fluorescent marker joined to the gene of interest by a T2A linker, and a plasmid backbone containing an MC1-DTA marker to select against random integration. sgRNA oligos were cloned into pX330 (Addgene #42230), which also expresses Cas9. Oligo sequences are provided in Supplementary Table 4. All plasmids will be made available on Addgene following acceptance of this paper.

Knock-in electroporations were performed as previously described (*25*). In summary, 1 µg donor plasmid and 1 µg sgRNA/Cas9 plasmid were co-electroporated into 200,000 hiPSCs using a Lonza Nucleofector with 20 µL P3 solution and pulse setting CA-137. Colonies were picked after selection and genotyped by PCR. Successful knockin, absence of oF-target edits, and euploidy of the reporter lines were all confirmed by whole genome sequencing (Novogene, 10X coverage) and SeqVerify computational analysis (*18*). Low-passage cells were cryopreserved using CryoStor CS10 (Stemcell Technologies) and banked for future use.

Functional validation of the reporter alleles was performed by CRISPRa. For each allele, three CRISPRa plasmids were constructed, each containing a sgRNA targeting the promoter as well as a doxycycline-inducible expression cassette for dCas9-VPR (*28*). Equimolar mixtures of plasmids (1 µg total) were electroporated into 200,000 hiPSCs using a Lonza Nucleofector with 20 µL P3 solution and pulse setting CA-137. After two days, cells were harvested with Accutase and analyzed by flow cytometry (Fig. S1).

### Identification of candidate meiosis-promoting factors

We obtained a human fetal germ cell scRNAseq dataset from Garcia-Alonso *et al.* 2022 (https://www.reproductivecellatlas.org/gonads.html) (*16*). We performed pySCENIC analysis (*29*), which involves inferring a gene regulatory network, finding transcription factor (TF) regulons based on known binding motifs, and ranking regulon activity in each cell type. We chose 21 TFs to screen based on high activity in STRA8+ and/or meiotic oogonia. Because regulon analysis ignores non-TF factors, including RNA-binding proteins, we selected 18 additional factors to screen based on a gene regulatory network analysis. In this analysis, we found differentially expressed genes between meiotic oogonia and all other cell types, then calculated which genes in the regulatory network were upstream of the differentially expressed genes, multiplying network edge weights by differential expression fold-changes to calculate a weighted score. We included a further 12 factors based on literature reports of pro-meiotic function. Finally, we included 27 factors from the Cancer Pathways ORFs library (*30*), which contains modulators of several common cellular signaling pathways. Our initial library for barcode enrichment screening contained 78 factors. For scRNAseq screening, we included an additional 10 factors involved in epigenetics and signal transduction, bringing the total to 88. A full list of the factors in our library is provided in Supplementary Table 2.

### Plasmid library construction and PiggyBac transposon integration

Expression plasmids for 88 candidate regulatory factors were constructed by MegaGate cloning (*31*) into a barcoded PiggyBac destination vector containing a doxycycline-inducible promoter (*28*), as well as a puromycin selection marker. Barcodes and transgene sequences were verified by Sanger sequencing. For later experiments, an alternative version of the plasmid lacking the barcode and containing a hygromycin selection marker was constructed for top candidate factors. For constitutive BCL2 expression, a version of the plasmid was constructed with an EF1a promoter instead of a doxycycline-inducible promoter. All plasmids will be made available on Addgene following acceptance of this paper.

Plasmids were pooled and co-electroporated into iPSCs along with a PiggyBac transposase expression plasmid (Systems Bioscience), using a Lonza Nucleofector with pulse setting CA-137. For medium-copy (3–5 per cell) integration, 5 fmol of pooled library and 500 ng of transposase plasmid were used per 200,000 hiPSCs and 20 µL of P3. For high-copy (10–50 per cell) integration, 50 fmol of pooled library was used instead. Average integration numbers were previously characterized for the same transposons used in this study (*28*), but not evaluated directly. Selection was performed with puromycin (400 ng/mL) and/or hygromycin (50 µg/mL) beginning 2 days after nucleofection and continuing for at least 5 additional days before subsequent experiments.

### Flow cytometry

Flow cytometry was performed on a BD LSR Fortessa instrument. Cell sorting was performed on a Sony SH800S sorter using a 100 µm chip. Compensation controls were acquired using cells in which mGreenLantern and tdTomato reporters had been activated by CRISPRa. DAPI (100 ng/mL) was used to stain dead cells for exclusion. Data analysis was performed using the Cytoflow python package (v. 1.0.0) (*32*). Representative gating is shown in Fig. S3A.

### Immunofluorescence microscopy

Cells were cultured on Matrigel-coated ibidi dishes (8-well, cat# 80826; or 96-well, cat# 89626). For 8-well dishes, 100 µL of staining solutions and 200 µL of wash solutions were used per well. For 96-well dishes, 50 µL of staining solutions and 100 µL of wash solutions were used per well. All steps except primary antibody incubation were performed at room temperature.

Cells were washed with PBS and fixed with 4% PFA in PBS for 10 minutes, followed by one 5-minute wash with PBS and one 5-minute wash with PBST (0.1% Triton X-100 in PBS). Cells were incubated with blocking solution (1% BSA and 5% normal donkey serum in PBST) for 20-30 minutes, followed by an overnight incubation at 4 °C with primary antibodies in blocking solution. Three 5-minute PBST washes were performed, followed by a 1-hour incubation with secondary antibodies and DAPI (1 µg/mL) in blocking solution. Two 5-minute PBST washes were performed, and the cells were stored in the dark at 4°C in PBST prior to imaging on a Zeiss LSM980 confocal microscope using a LD C-Apochromat 40x/1.1 water immersion objective. A list of antibodies used, and their dilutions, is provided in Supplementary Table 5.

Initial image processing was performed using FIJI (ImageJ version 2.14.0/1.54f) (*33*). Brightness and contrast were adjusted equally across all images from each experiment. Segmentation and quantification were performed using the cellpose Python package (version 2.2.3) (*34*). Thresholding was performed by using a negative control (typically hiPSCs) to determine the average background staining intensity, subtracting this average background intensity from all regions, then classifying any region that was >2x brighter than then 95^th^ percentile of the negative control as positive. Analysis code is available on Github: https://github.com/mpiersonsmela/meiosis

### Barcode enrichment screening

Reporter iPSCs containing integrated expression transposon vectors were harvested with Accutase. Cells were seeded on Matrigel-coated 6-well plates (500,000 cells per plate) in mTeSR1 medium (Stemcell Technologies) with doxycycline (1 µg/mL) and Y-27632 (10 µM). A media change was performed after 24 hours. At this point, candidate differentiation media were tested, based on previous claims of meiosis induction in the literature. These included: A-MEM with 1X Glutamax, 1X Insulin-Transferrin-Selenium-X supplement, 0.2% BSA, 0.2% chemically defined lipid concentrate, 200 µg/mL ascorbic acid, 1 ng/mL FGF2, and 20 ng/mL GDNF (*35*).

Nutrient restriction/retinoic acid: EBSS with 0.1X IMDM, 0.1X supplements (N2, glutamax, sodium pyruvate, MEM essential vitamins, non-essential amino acids), 0.1% FBS, 0.5% Knockout Serum Replacement, 0.6 mg/mL glucose, 0.1 mg/mL lactic acid, 0.5 mg/mL BSA, 5 µM 2-mercaptoethanol, 10 µM ascorbic acid, 1 µg/mL biotin, 3 ng/mL beta-estradiol, 1 ng/mL FGF2, 1.5 ng/mL GDNF, and 100 nM retinoic acid (*7*).

We additionally tested mTeSR1 (which maintains primed pluripotency), HENSM (which induces naïve pluripotency) (*36*), StemPro-34 based spermatogonial stem cell culture medium (*37*), and APEL2 (Stemcell Technologies; a “neutral” medium lacking growth factors).

The cells were cultured in these six media, all supplemented with 1 µg/mL doxycycline, for six additional days. A media change was performed each day. Cells were harvested with Accutase on day seven post-induction, and reporter-positive cells were isolated by FACS. DNA was extracted from REC8+, SYCP3+, and DDX4+ cells, as well as the pre-sorting population and pre-differentiation population. Barcodes were amplified by two rounds of PCR and sequenced on an Illumina MiSeq as previously described (*25*). Barcode enrichment was calculated by comparing the barcode frequencies in reporter-positive and pre-sorting cells. We also sequenced barcodes from the pre-differentiation cells, but did not use these for analysis since effects were dominated by changes in cell growth rate.

### CRISPRi of epigenetic factors

Guide RNAs targeting the promoters of ten epigenetic factors (*MAX, MGA, E2F6, RNF2, PCGF6, SETDB1, DNMT1, DNMT3A, DNMT3B,* and *UHRF1*; three guides per gene) were cloned into a doxycycline-inducible dCas9-KRAB expression transposon plasmid (*28*). Each transposon plasmid was integrated into SYCP3 reporter iPSCs as described above. iPSCs were treated with 1 µg/mL doxycycline in mTeSR1 for six days, harvested with Accutase, and analyzed by flow cytometry (Fig. S4). Additionally, knockdown efficiency was evaluated by qPCR with PowerUp SYBR Green Master Mix (Thermo Fisher) using *GAPDH* as a control gene (Fig. S4C). Primers used are given in Supplementary Table 4.

### Screening of conditions for reporter activation

Several flow cytometry experiments were performed in order to optimize conditions for activating REC8 and SYCP3 expression in male (PGP1) and female (F3) reporter lines. First, expression vectors for sixteen promising factors (Fig. S3) chosen based on barcode enrichment data were individually integrated into REC8 and SYCP3 reporter iPSC lines, and expression was induced by treatment with 1 µg/mL doxycycline in APEL2 medium following the protocol described above in the barcode enrichment section.

Second, a fractional factorial screen for REC8 activation was conducted in F3 and PGP1 D4TR8G reporter lines, using 32 combinations of seven promising factors and following the same differentiation protocol. (Fig. 1C)

Third, REC8, SYCP3, and DDX4 activation was measured after performing differentiation using expression of STRA8, HOXB5, and BCL2 in the presence of different basal media and additives. Four basal media were tested: mTeSR1 (Stemcell Technologies); APEL2 (Stemcell Technologies); DMEM/F12 with 1X Glutamax and 10% KSR (Gibco); and Advanced RPMI with 1X nonessential amino acids, 1X Glutamax, and 0.5X B27 supplement minus Vitamin A (all Gibco). For each of the differentiation media, five additives were tested, as well as a doxycycline-only control. The additives and their concentrations were: 1 µM retinoic acid, 1 µM AM580, 25 µM PRT4165, 20 µM RB3, and 1 mM sodium valproate. A media change was performed every day. At days 6, 7, and 8 post-induction, cells were harvested with Accutase and analyzed by flow cytometry (Fig. 1D)

Fourth, REC8, SYCP3, and DDX4 activation was measured in cells treated with or without DNMT1 inhibitor (5 µM GSK3484862). Differentiation was performed using expression of STRA8, HOXB5, and BCL2 in APEL2 medium supplemented with 1 µM AM580 and 1 µg/mL doxycycline. Cells were harvested with Accutase after 7 days and analyzed by flow cytometry (Fig. 1E).

### scRNAseq screening

Prior to scRNAseq, the following six cell populations were generated by integration of transposon expression vectors:

- PGP1 D4TR8G, with BCL2, HOXB5, and STRA8 under hygromycin selection
- F3 D4TS3G, with BCL2, HOXB5, and STRA8 under hygromycin selection
- PGP1 D4TR8G, with BCL2, HOXB5, and STRA8 under hygromycin selection and the full pool of 88 candidate factors under puromycin selection
- F3 D4TS3G, with BCL2, HOXB5, and STRA8 under hygromycin selection and the full pool of 88 candidate factors under puromycin selection
- PGP1 D4TR8G, with the full pool of 88 candidate factors under puromycin selection
- F3 D4TS3G, with the full pool of 88 candidate factors under puromycin selection

The cells were differentiated according to the following method, which had been chosen based on its ability to activate REC8 and SYCP3 expression. Cells containing integrated expression vectors were seeded in mTeSR1 containing 10 µM Y-27632, 5 µM GSK3484862, and 1 µg/mL doxycycline. After 24 hours, the medium was changed to APEL2 with 5 µM GSK3484862, 1 µM AM580, and 1 µg/mL doxycycline. A media change was performed every other day. After a total of seven days of differentiation, cells were harvested with Accutase and sorted based on reporter expression. Cells were fixed using a Parse fixation kit, and scRNAseq library preparation was performed using a Parse WT Mega v2 kit. A list and description of samples is provided in Supplementary Table 6. Sequencing was performed on an Illumina Novaseq X Plus using three full 10B PE150 flowcells. Alignment and counts matrix generation was performed using the Parse Biosciences pipeline (v.0.9.6).

To enrich the library for barcode sequences, we first performed PCR with biotinylated primers to generate a dsDNA biotinylated bait containing the 120bp of sequence immediately 3’ of the barcode sequence in our expression vector. We isolated the bait DNA from the PCR using a 3X volume of ProNex beads, and eluted in 10 mM Tris pH 8.0 buffer.

Next, we used 200 fmol of bait DNA as a custom probe in the Parse Gene Capture kit, following the manufacturer’s protocol aside from this substitution. After qPCR to verify the barcode enrichment, the resulting library was sequenced on one lane of a NovaSeq X Plus 10B PE150 flowcell.

scRNAseq data were filtered by number of reads per cell (<100,000), number of genes detected (>1,000 and <14,000), and mitochondrial read percentage (<10%). Transgene barcode reads were merged into the dataset by matching cell barcodes. Analysis was performed using scanpy (*38*) for normalization, integration with the fetal germ cell reference atlas, and gene scoring (*16*).

### Refinement of factors for meiosis induction

Several experiments were performed in order to optimize the protocol for meiosis induction. First, for each of the 23 candidate factors identified by the scRNAseq screen (excluding HOXB5, which was already integrated), an expression vector with a puromycin selection marker was integrated into PGP1 D4TR8G and F3 D4TS3G reporter hiPSCs which already contained expression vectors for BCL2, HOXB5, and STRA8 with hygromycin selection markers. Two control conditions were additionally included: BCL2, HOXB5, and STRA8 only; and a no-factor control. Cells were differentiated using the same conditions as for the scRNAseq experiment (APEL2 with doxycycline, AM580, and GSK3484862). Cells were fixed and stained for SYCP3, HORMAD1, and DDX4 after 7 days of differentiation.

Second, expression vectors for the seven top factors identified in the previous experiment (BCL2, HOXB5, STRA8, myr-AKT1, BOLL, MEIOC, and MEIOSIN) were pooled and integrated into PGP1 D4TR8G and F2 D4TDZG hiPSCs. Cells were differentiated in the same manner, and fixation and staining for SYCP3, HORMAD1, and γH2AX was performed after 7, 9, 13, and 16 days of differentiation. As a control, the same differentiation and staining was performed using only BCL2, HOXB5, and STRA8.

Third, a fractional factorial screen was performed in order to identify the contributions of the seven top factors. Sixteen combinations of the seven factors (including one control combination lacking all factors) were integrated into F2 D4TDZG, F3 D4TS3G, and PGP1 D4TR8G reporter lines. Additionally, expression vectors for DAZL and BOLL were integrated in order to evaluate previous claims that those factors alone could induce meiosis (*11*, *14*). Cells were fixed and stained for SYCP3, HORMAD1, and γH2AX after 13 days of differentiation. Additional cells were analyzed by flow cytometry for reporter expression.

Fourth, a full factorial screen was performed in order to confirm the best factors for meiosis induction. A constitutive EF1a-driven BCL2 expression plasmid was integrated into F2 D4TDZG, F3 D4TT2G, and PGP1 D4TR8G reporter hiPSCs under hygromycin selection. Then, all eight possible combinations of HOXB5, BOLL, and MEIOC expression vectors were integrated under puromycin selection. Cells were fixed and stained for after 13 days of differentiation, with the first 3 days at 37 °C and the remainder at 34 °C. Two stains were used: SYCP3, HORMAD1, and γH2AX; and TEX12 and SYCP3.

### Timing of media additives and factor expression

A Shield1-inducible HOXB5 PiggyBac transposon plasmid was constructed using an EF1a promoter driving HOXB5 with a C-terminal degradation domain, which could be stabilized by addition of Shield1. This plasmid, along with expression plasmids for constitutive EF1a-driven BCL2 and doxycycline-inducible BOLL, were integrated into F2 D4TDZG, F3 D4TS3G, and PGP1 D4TR8G reporter lines. Cells were seeded at a density of 50,000/cm^2^ in Matrigel coated 96-well ibidi plates in mTESR1 + 10 µM Y-27632. After 24 hours, the medium was changed to APEL2. Subsequently, a full media change was performed every 48 hours, and cells were fixed and stained after 11 days of differentiation. This experiment was performed at 37 °C.

The following media additive conditions were tested:

1. No additives (negative control)
2. Full, continuous dose of all additives (positive control): 5 µM GSK3484862 (DNMT1i), 1 µM AM580, 1 µg/mL doxycycline, 500 nM Shield1
3. No doxycycline, 100 nM Shield1, full dose of DNMTi and AM580
4. 0.1 µg/mL doxycycline, no Shield1, full dose of DNMTi and AM580
5. 0.1 µg/mL doxycycline, 100 nM Shield1, full dose of DNMTi and AM580
6. 0.1 µg/mL doxycycline, 500 nM Shield1, full dose of DNMTi, and AM580
7. 1 µg/mL doxycycline, 100 nM Shield1, full dose of DNMTi and AM580
8. Full dose of all additives except doxycycline
9. Doxycycline added only after day 1. Other additives at full dose.
10. Doxycycline added only after day 3. Other additives at full dose.
11. Doxycycline added only after day 5. Other additives at full dose.
12. Doxycycline added only after day 7. Other additives at full dose.
13. Doxycycline added only after day 9. Other additives at full dose.
14. Full dose of all additives except Shield1
15. Shield1 added only after day 1. Other additives at full dose.
16. Shield1 added only after day 3. Other additives at full dose.
17. Shield1 added only after day 5. Other additives at full dose.
18. Shield1 added only after day 7. Other additives at full dose.
19. Shield1 added only after day 9. Other additives at full dose.
20. Full dose of all additives except AM580
21. AM580 added only after day 1. Other additives at full dose.
22. AM580 added only after day 3. Other additives at full dose.
23. AM580 added only after day 5. Other additives at full dose.
24. AM580 added only after day 7. Other additives at full dose.
25. AM580 added only after day 9. Other additives at full dose.
26. Full dose of all additives except DNMTi
27. DNMTi withdrawn after day 5. Other additives at full dose.
28. DNMTi withdrawn after day 7. Other additives at full dose.
29. DNMTi withdrawn after day 9. Other additives at full dose.
30. Shield1 added only from days 0–5, doxycycline added only from days 5–11, other additives at full dose.
31. Shield1 added only from days 0–7, doxycycline added only from days 5–11, other additives at full dose.
32. Shield1 added only from days 0–7, doxycycline added only from days 7–11, other additives at full dose.

### Evaluation of temperatures and timings for meiosis induction

As an initial experiment to evaluate the effects of lower temperature on male meiosis, PGP1 D4TR8G reporter hiPSCs containing integrated expression vectors for BCL2, HOXB5, BOLL, and MEIOC were seeded in 8-well ibidi dishes at 50,000 cells/cm2 in mTeSR1 with 5 µM GSK3484862, 1 µg/mL doxycycline, and 10 µM Y-27632. After 24 hours, the media was replaced with APEL2 containing 5 µM GSK3484862 and 1 µg/mL doxycycline. A 50% media change was performed every 2 days, and GSK3484862 was withdrawn starting on day 7. Initially, all cells were cultured at 37 °C. One plate was moved to a 34 °C incubator after the first day. Cells were fixed on day 13, stained for HORMAD1, SYCP3, and γH2AX, and imaged.

As a confirmatory experiment, F2 D4TDZG, F3 D4TS3G, and PGP1 D4TR8G reporter hiPSCs containing integrated expression vectors for BCL2, HOXB5, BOLL, and MEIOC were differentiated according to the same protocol. The following conditions were tested:

1. 34 °C starting on day 3, fixation at day 13
2. 34 °C starting on day 1, fixation at day 14
3. 34 °C starting on day 3, fixation at day 15
4. 34 °C starting on day 3, fixation at day 16
5. Continuous 37 °C, fixation at day 16
6. 34 °C starting on day 1, fixation at day 17
7. 34 °C starting on day 3, fixation at day 19
8. 34 °C starting on day 3, fixation at day 21

After fixation, cells were stained for HORMAD1, SYCP3, γH2AX, and actin, and imaged.

### Final protocol for meiosis induction

Based on the results of our screening and optimization experiments, which are described above, we have developed the following protocol for robust initiation of meiosis:

A constitutive or doxycycline-inducible expression vector for the anti-apoptotic factor BCL2, as well as doxycycline-inducible expression vectors for meiosis-promoting factors (HOXB5, BOLL, and/or MEIOC), are integrated into human iPSCs using PiggyBac transposase. The iPSCs are seeded at 50,000 cells/cm^2^ on Matrigel-coated plates in mTeSR1 supplemented with 5 µM GSK3484862, 1 µg/mL doxycycline, and 10 µM Y-27632. For 6-well plates, 1.5 mL of medium is used per well, and volumes for smaller plates are scaled down proportionally to their surface area. After one day, the media is replaced with APEL2 containing 5 µM GSK3484862 and 1 µg/mL doxycycline. A 50% media change is performed every 2 days, and GSK3484862 is withdrawn starting on day 7. Initiation of meiosis is complete by roughly day 13.

### Timecourse scRNAseq and imaging

Plasmids for constitutive expression of BCL2 and doxycycline-inducible expression of HOXB5, BOLL, and MEIOC were integrated into F2 D4TDZG, F3 D4TT2G, and PGP1 D4TR8G reporter hiPSCs. Meiosis was initiated following the final optimized protocol, with cells cultured on 8-well ibidi dishes for immunofluorescence imaging and 12-well plates for scRNAseq. At each day from day 0 (hiPSC) to day 15, cells were fixed for imaging and harvested for scRNAseq. Stains used for imaging were: rabbit anti-HORMAD1, goat anti-SYCP3, mouse anti γH2AX, and rat anti-T2A; and rabbit anti-TEX12, goat anti-SYCP3, mouse anti-RAD51, and rat anti-KI67. Samples for scRNAseq were counted and fixed using the Parse Biosciences fixation kit. Library preparation was performed using the Parse Biosciences WT v3 kit. Sequencing was performed on two lanes of a Novaseq X Plus 25B PE150 flowcell. Alignment and counts matrix generation was performed using the Parse Biosciences pipeline (v.1.2.1). Filtering, normalization, and cell type annotation were performed in scanpy as described above. To build a reference atlas, we combined the fetal gonad atlas and adult testis atlas (*16*, *22*), removing somatic cell types from the testis atlas and only keeping genes that were expressed in both atlases.

### Statistics and data analysis

Results of fractional factorial screens were analyzed by fitting linear models using the lm function in R (version 4.3.2). For flow cytometry data, values expressed as a proportion (0 - 100%) were logit-transformed before fitting the model. Significance calculations in bar plots were performed using two-tailed Mann-Whitney U tests.

---

**Fig. S1.**
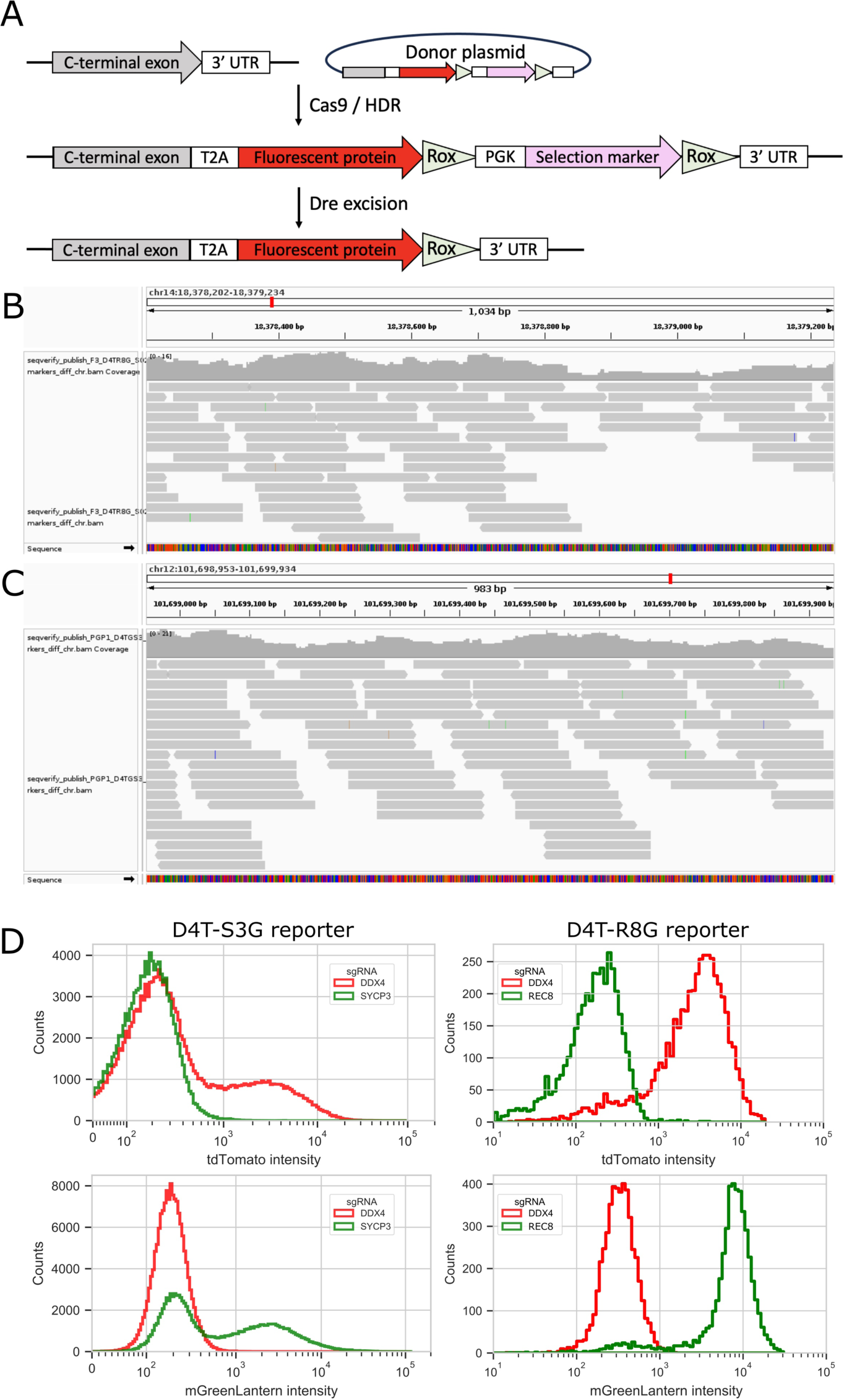
Construction and validation of hiPSC reporter lines. (**A**) Knock-in editing strategy. (**B**) SeqVerify validation of REC8 reporter allele using whole genome sequencing. (**C**) SeqVerify validation of SYCP3 reporter allele using whole genome sequencing. (**D**) Functional validation of reporter hiPSCs using flow cytometry and CRISPRa with gRNAs targeting the promoters of *REC8, SYCP3*, and *DDX4*.

**Fig. S2.**
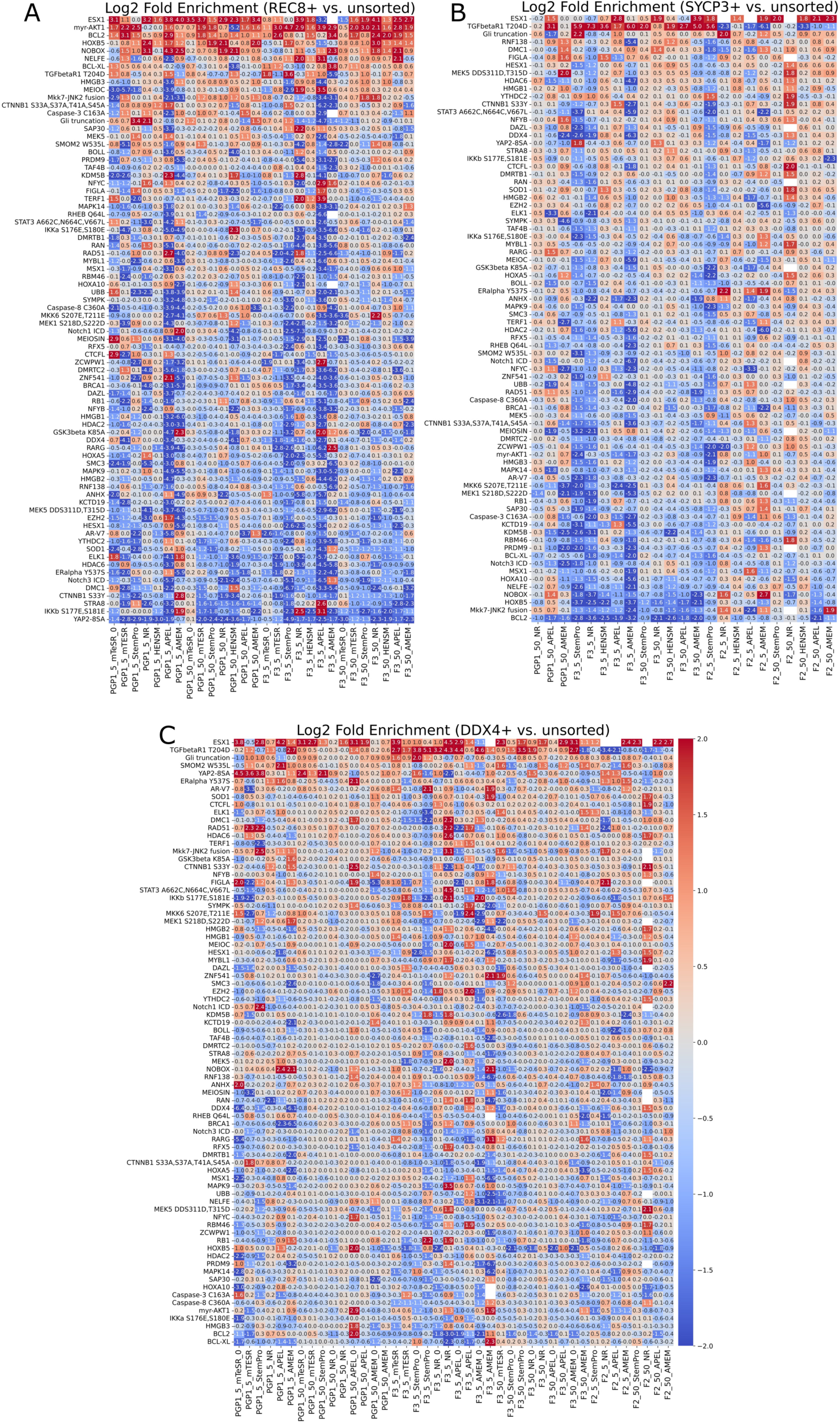
Barcode enrichment results. for (**A**) REC8, (**B**) SYCP3, and (**C**) DDX4. Reporter hiPSCs (F2, F3, and PGP1) were nucleofected with low (5 fmol) or high (50 fmol) doses of plasmid library pool, treated with doxycycline to induce expression, and differentiated in various media (mTeSR, StemPro, nutrient restriction (NR), HENSM, spermatogonial stem cell medium (AMEM), and APEL2). Reporter-positive cells were sorted after 7 days, and barcode frequencies were compared to unsorted cells.

**Fig. S3.**
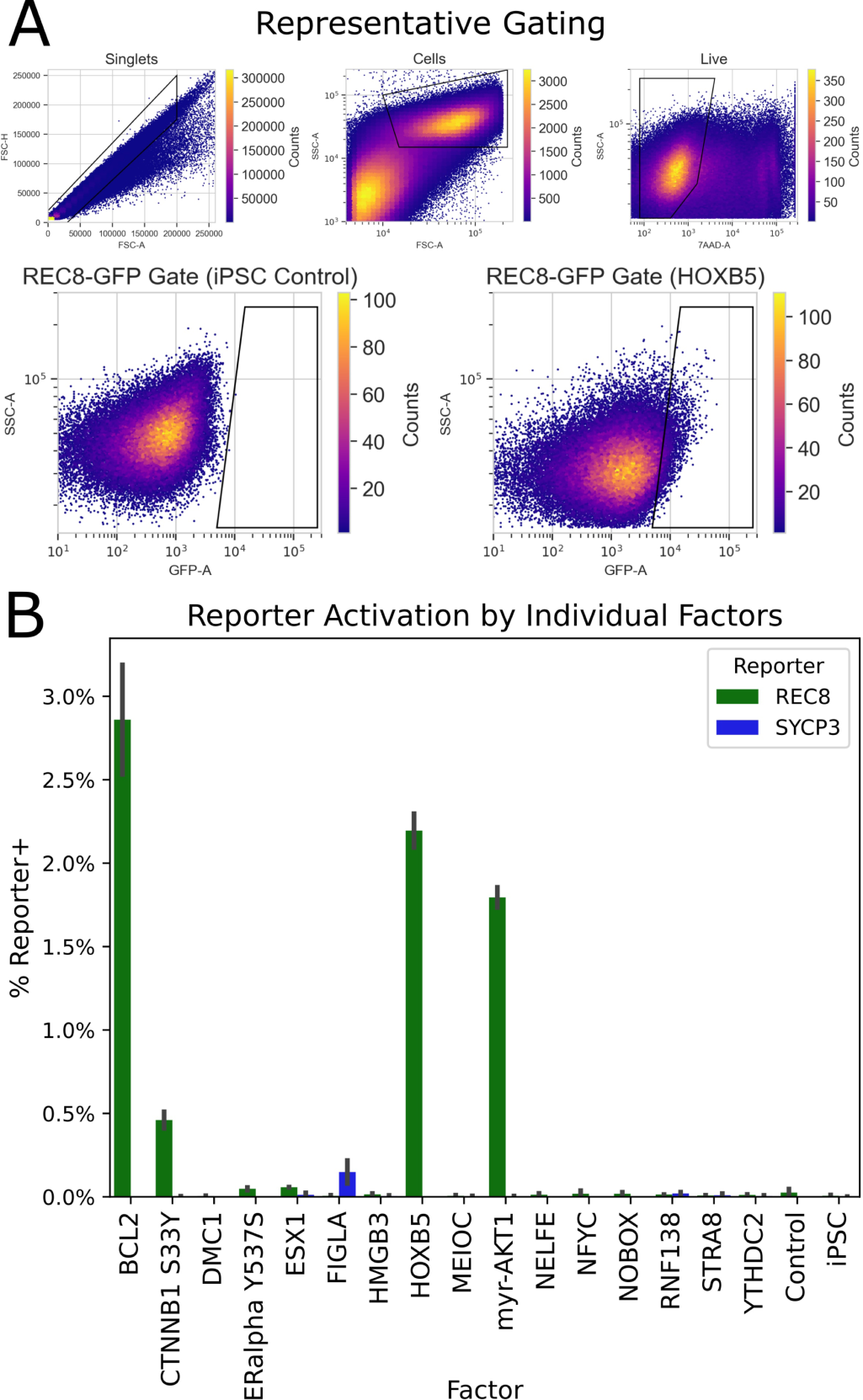
Flow cytometry analysis of reporter expression induced by individual factors. (A) Representative gating strategy for singlets, cells, live cells, and reporter-positive cells. (B) Activation of REC8 and SYCP3 reporters by sixteen individual factors chosen based on barcode enrichment results (n = 2 biological replicates per factor per reporterX). Error bars are standard error of the mean.

**Fig. S4.**
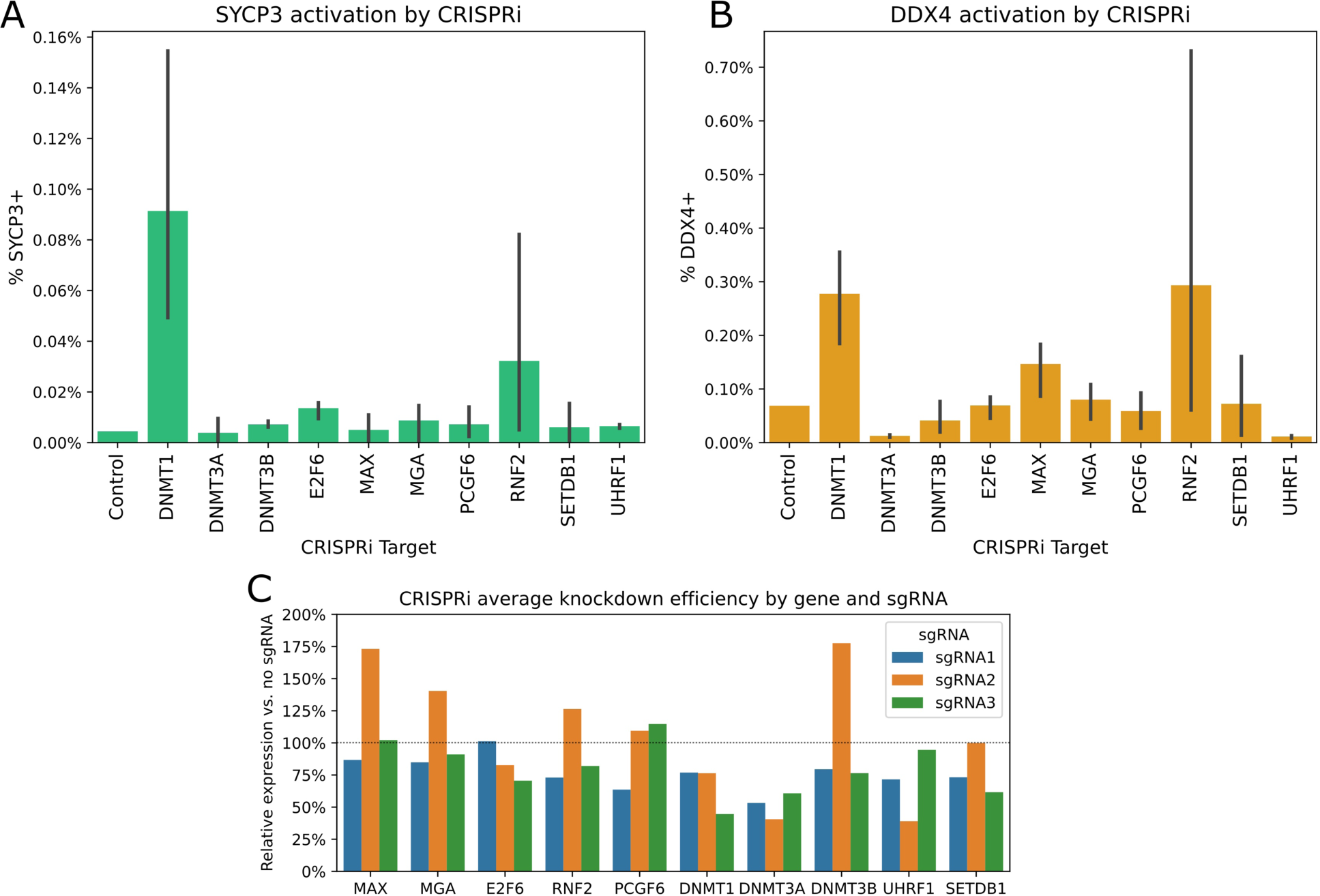
Pilot screen for activation of SYCP3 and DDX4 expression upon CRISPRi knockdown of ten epigenetic modifiers. (**A**) Activation of SYCP3 expression measured by flow cytometry (n = 3 sgRNAs per gene). (**B**) Activation of DDX4 expression measured by flow cytometry (n = 3 sgRNAs per gene). (**C**) qPCR measurement of average knockdown efficiency (n = 2 technical replicates per guide), calculated by 2^-ΔΔCt^ with *GAPDH* as a reference gene. The bulk knockdown efficiency was poor for most guides, although it is possible that subpopulations of cells experienced a greater knockdown.

**Fig. S5.**
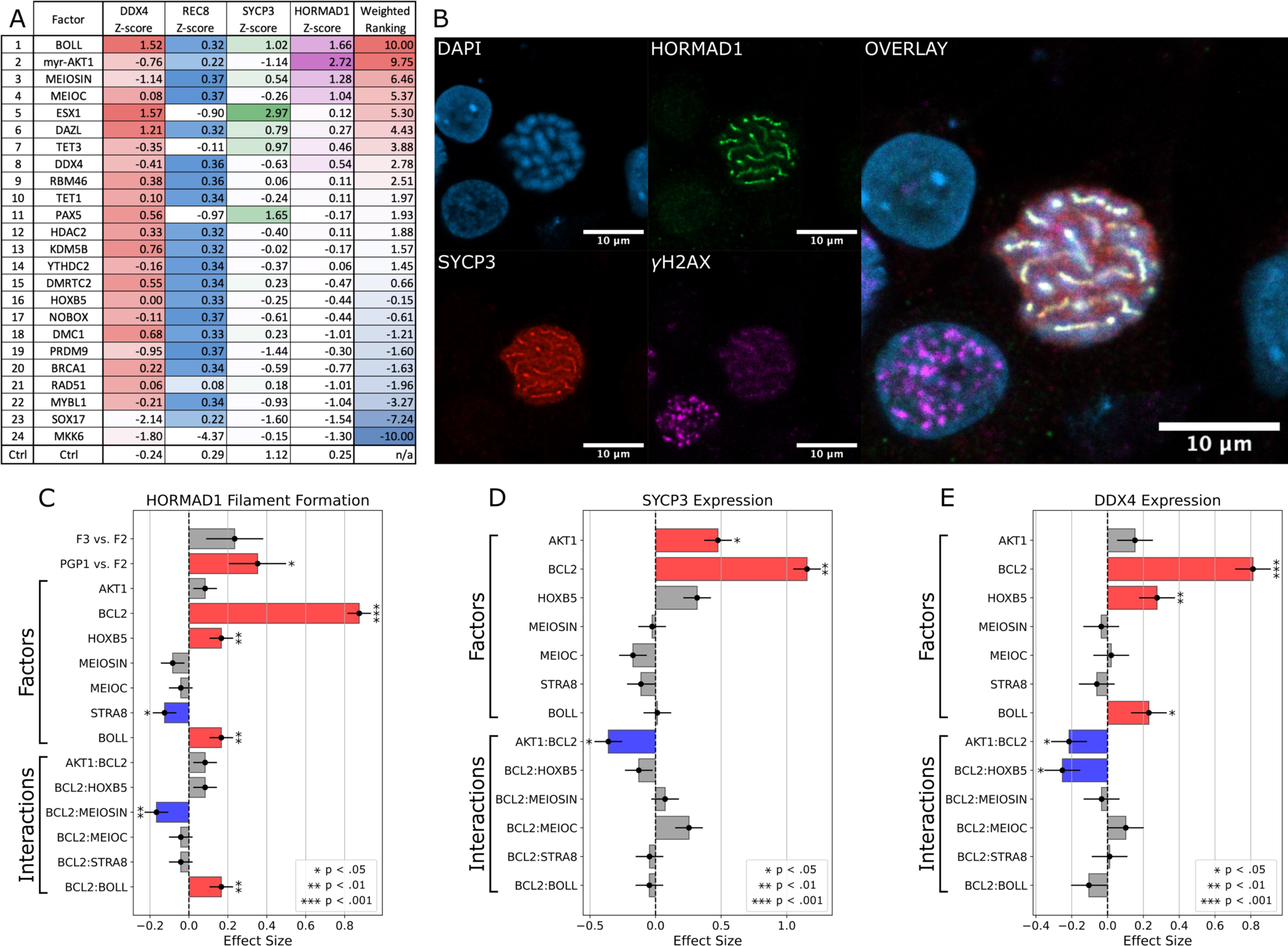
Optimization of factors for meiosis induction. (**A**) 24 candidate factors identified by an scRNAseq screen (Fig. 2D) were each co-expressed with STRA8, BCL2, and HOXB5. Reporter expression was analyzed by flow cytometry, and HORMAD1 expression was analyzed by immunofluorescence microscopy. Factors were ranked according to the results. (**B**) HORMAD1 and SYCP3 filament formation observed by immunofluorescence microscopy after twelve days of expression of seven top factors (STRA8, BCL2, HOXB5, BOLL, AKT1, MEIOSIN, and MEIOC). (**C**) Results of a fractional factorial screen of the seven top factors, for HORMAD1 filament formation. (**D**) Results for SYCP3 expression. (**E**) Results for DDX4 expression.

**Fig. S6.**
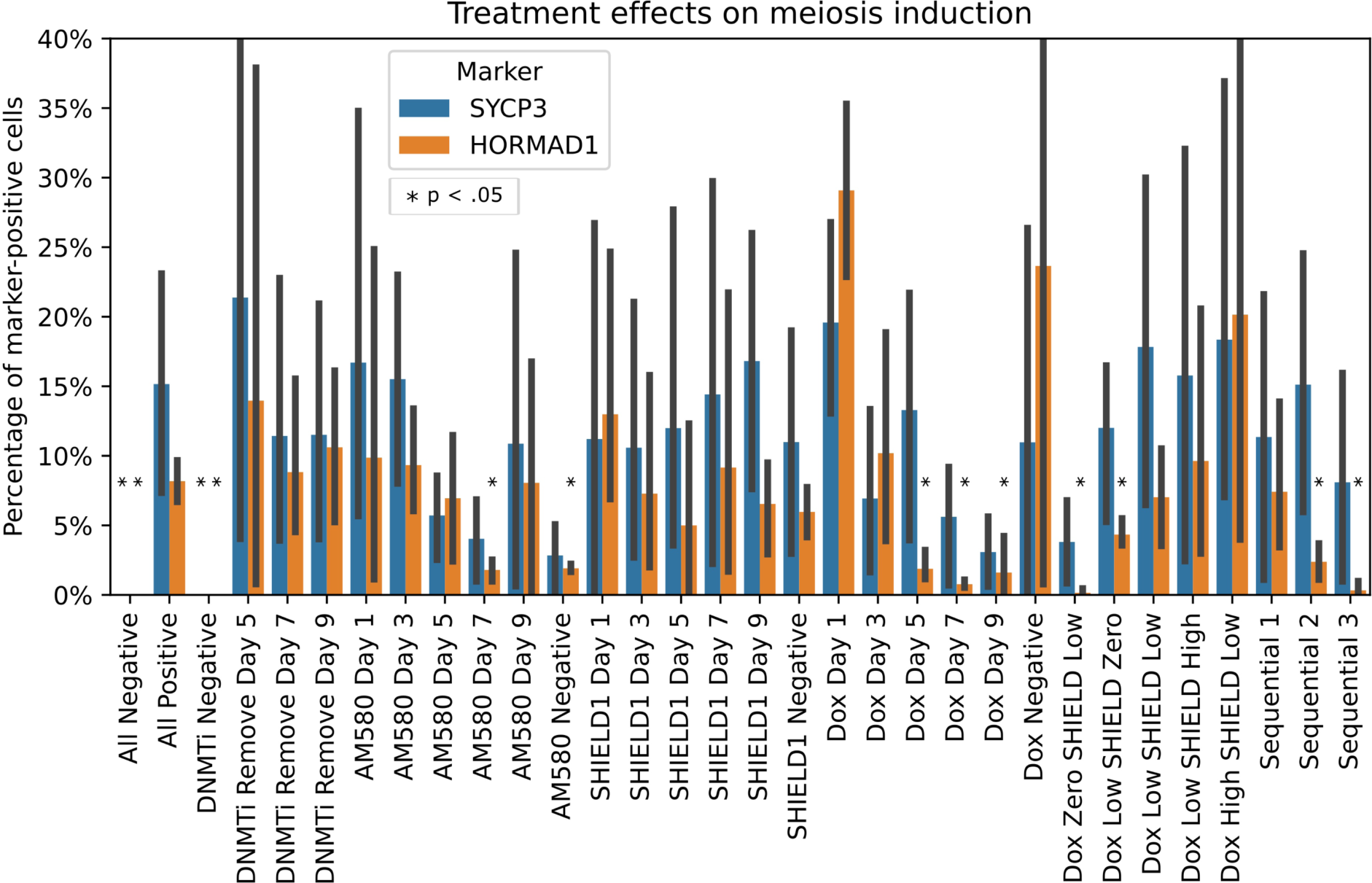
Quantification of the eGects of omitting various components of the meiosis induction protocol, or reducing their doses. Significance test comparisons are to the “all positive” control.

**Fig. S7.**
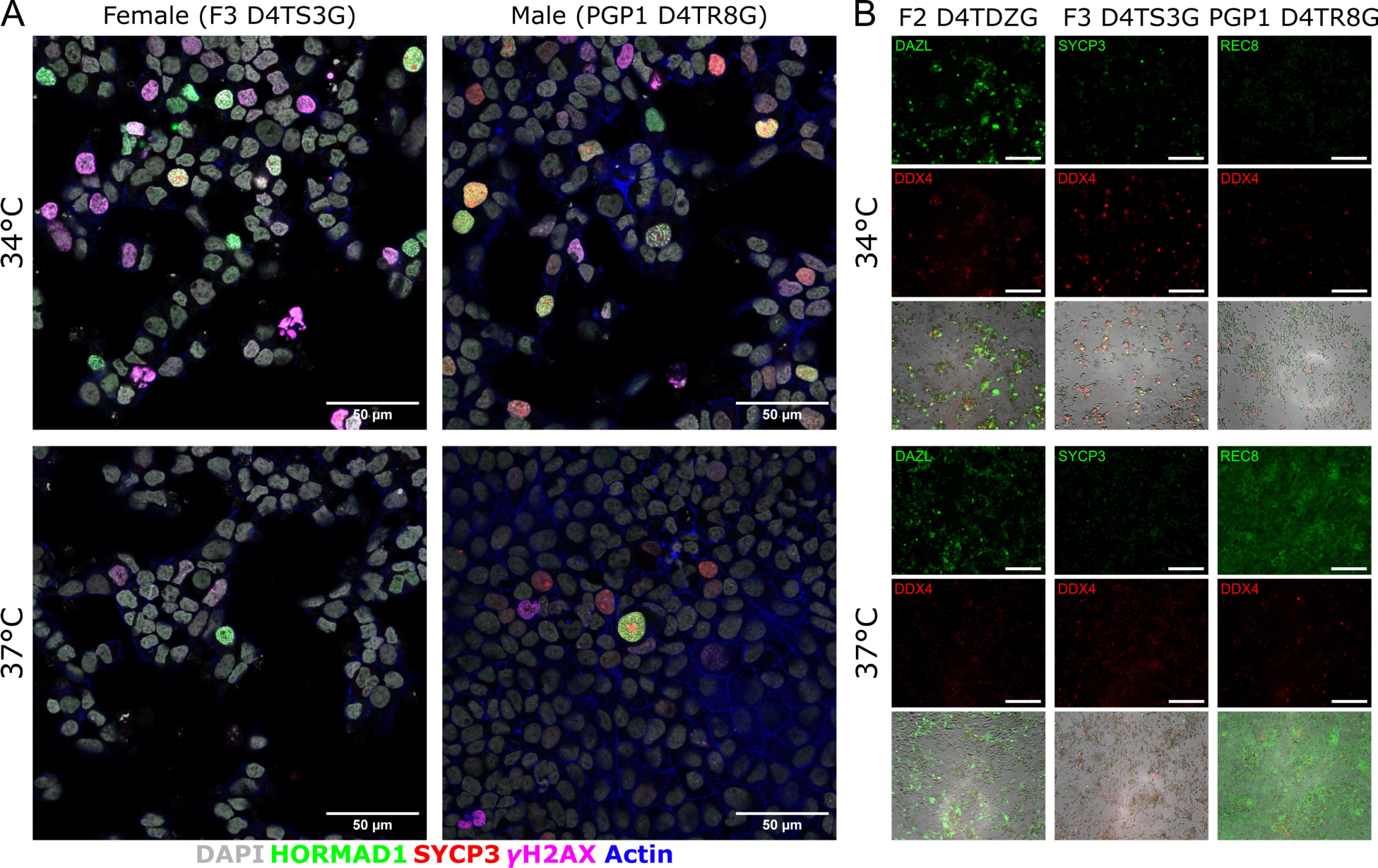
Microscope images of meiosis induction from male and female hiPSCs at 34 °C vs. 37 °C. (**A**) Immunofluorescence microscopy, staining for DNA (DAPI; gray), HORMAD1 (green), SYCP3 (red), γH2AX (magenta), and actin (phalloidin; blue). Scale bar is 50 µm. (**B**) Live imaging of fluorescent reporter expression of DAZL, SYCP3, REC8, and DDX4. Scale bar is 200 µm.

**Fig. S8.**
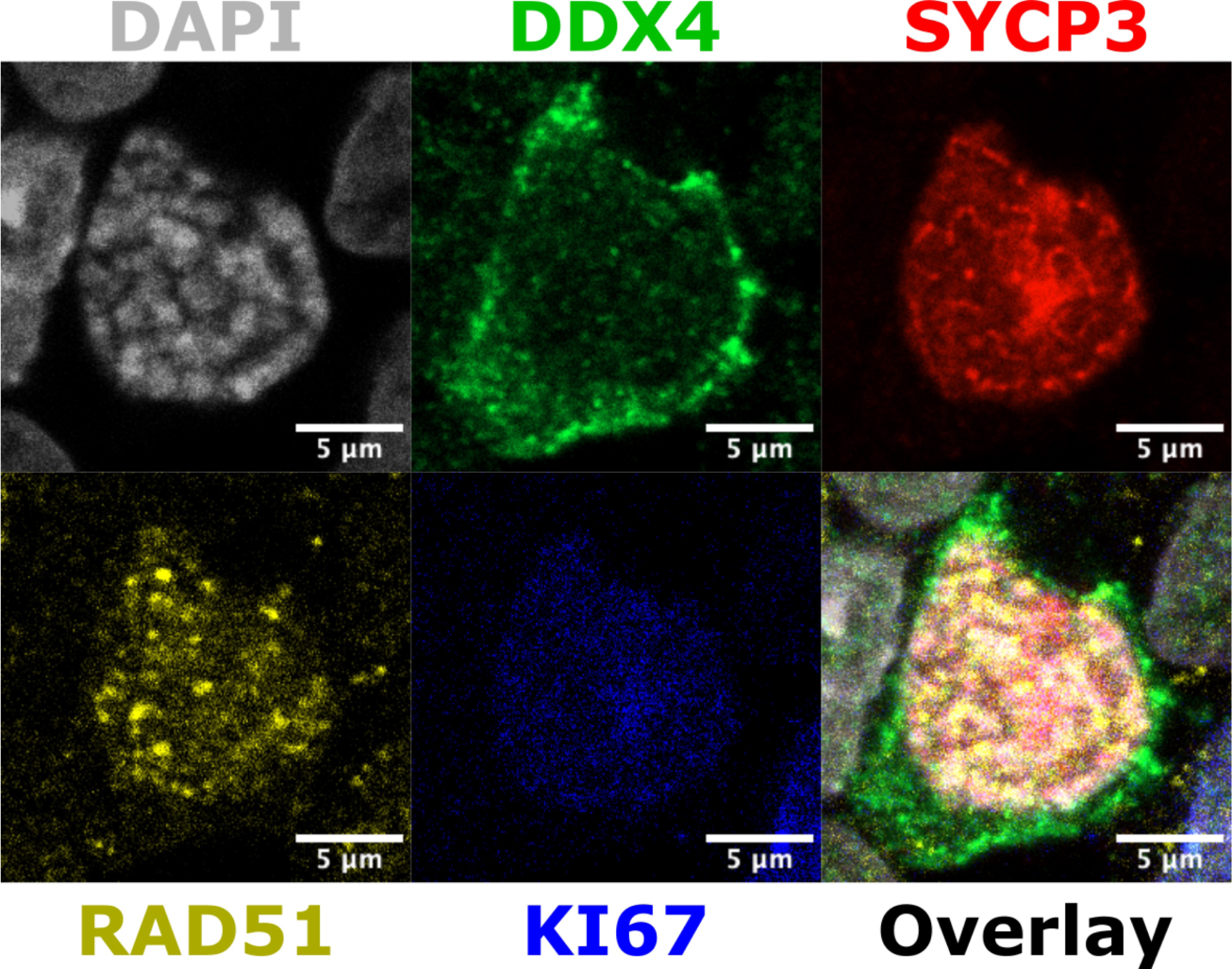
Immuno-staining for DDX4, SYCP3, RAD51, and KI67 in a day 15 female meiotic cell. Cytoplasmic DDX4 staining, filamentous SYCP3 staining, and nuclear RAD51 foci are observed.

**Fig. S9.**
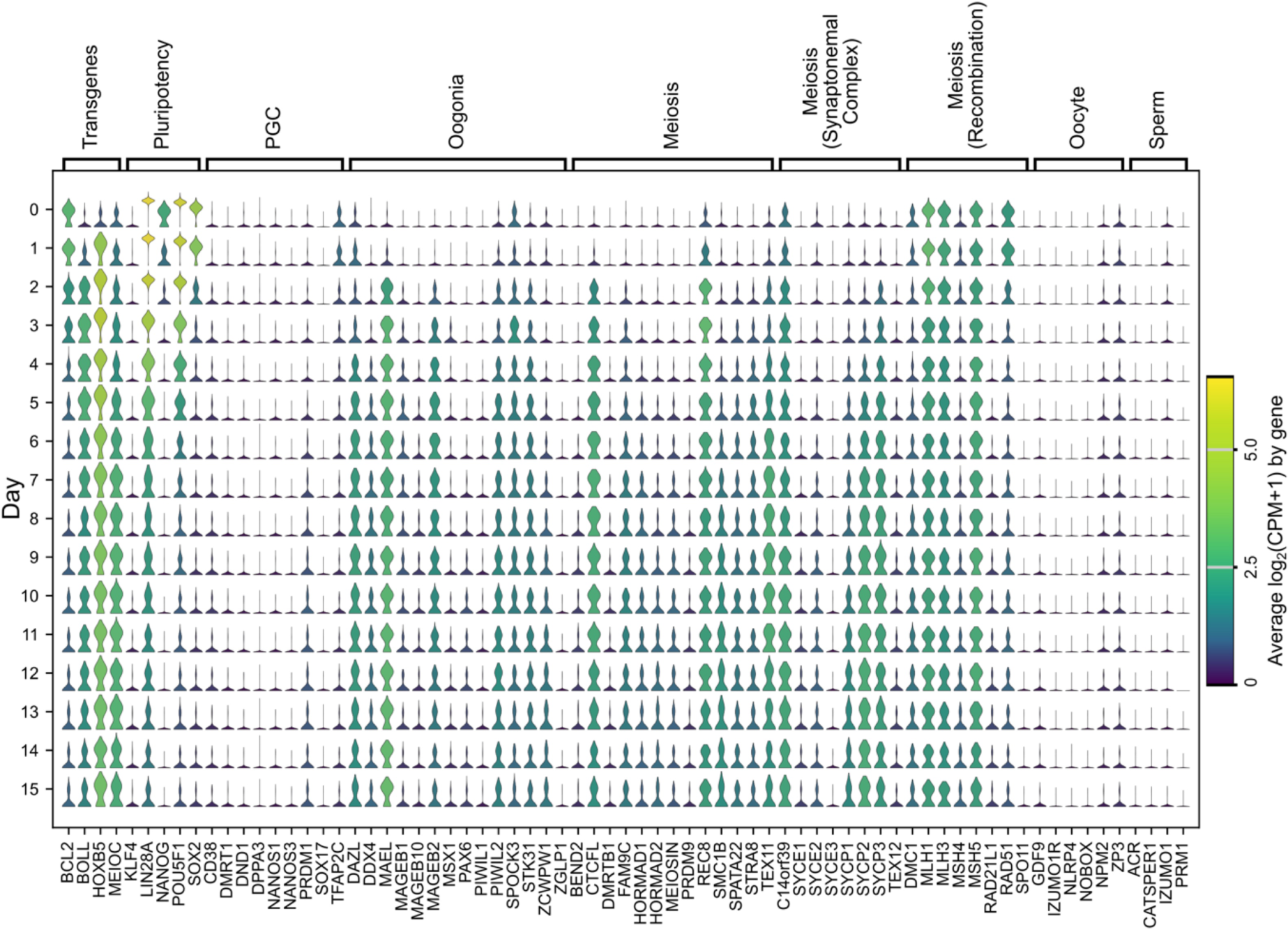
Violin plots showing expression of marker genes from days 0 to 15 of meiosis induction. Units for the color scale are log_2_(CPM+1). Categories include: exogenous transgenes, pluripotency markers, primordial germ cell markers, oogonia markers, meiosis markers, synaptonemal complex components, recombination markers, oocyte markers, and sperm markers.

**Supplementary Table 1.**
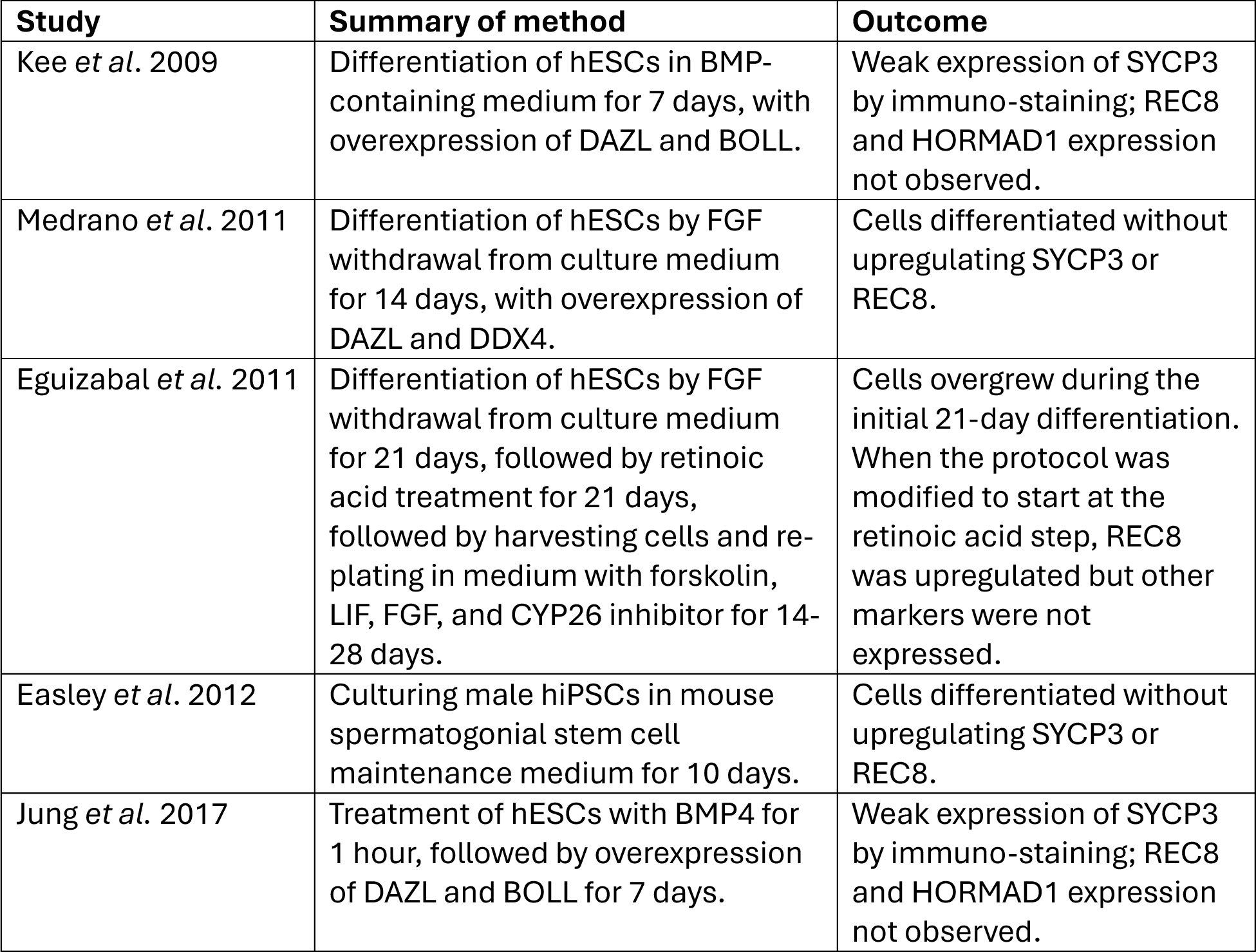
Replication attempts of previous reported methods for inducing meiosis from human pluripotent stem cells. The methods were carried out as described in their respective papers (see References), although PiggyBac transposon plasmids were used instead of lentivirus for transgene overexpression.

**Supplementary Table 2.**
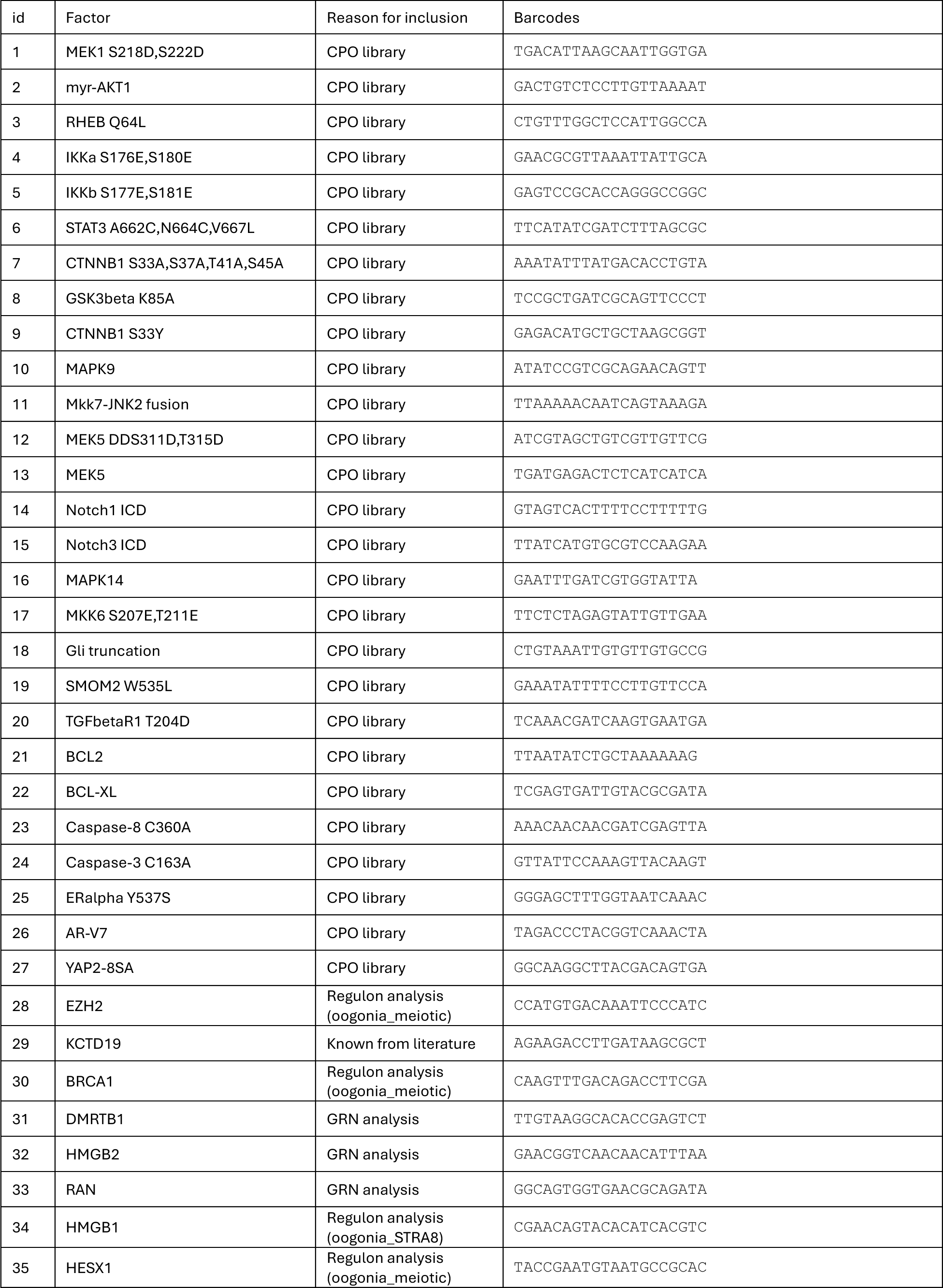
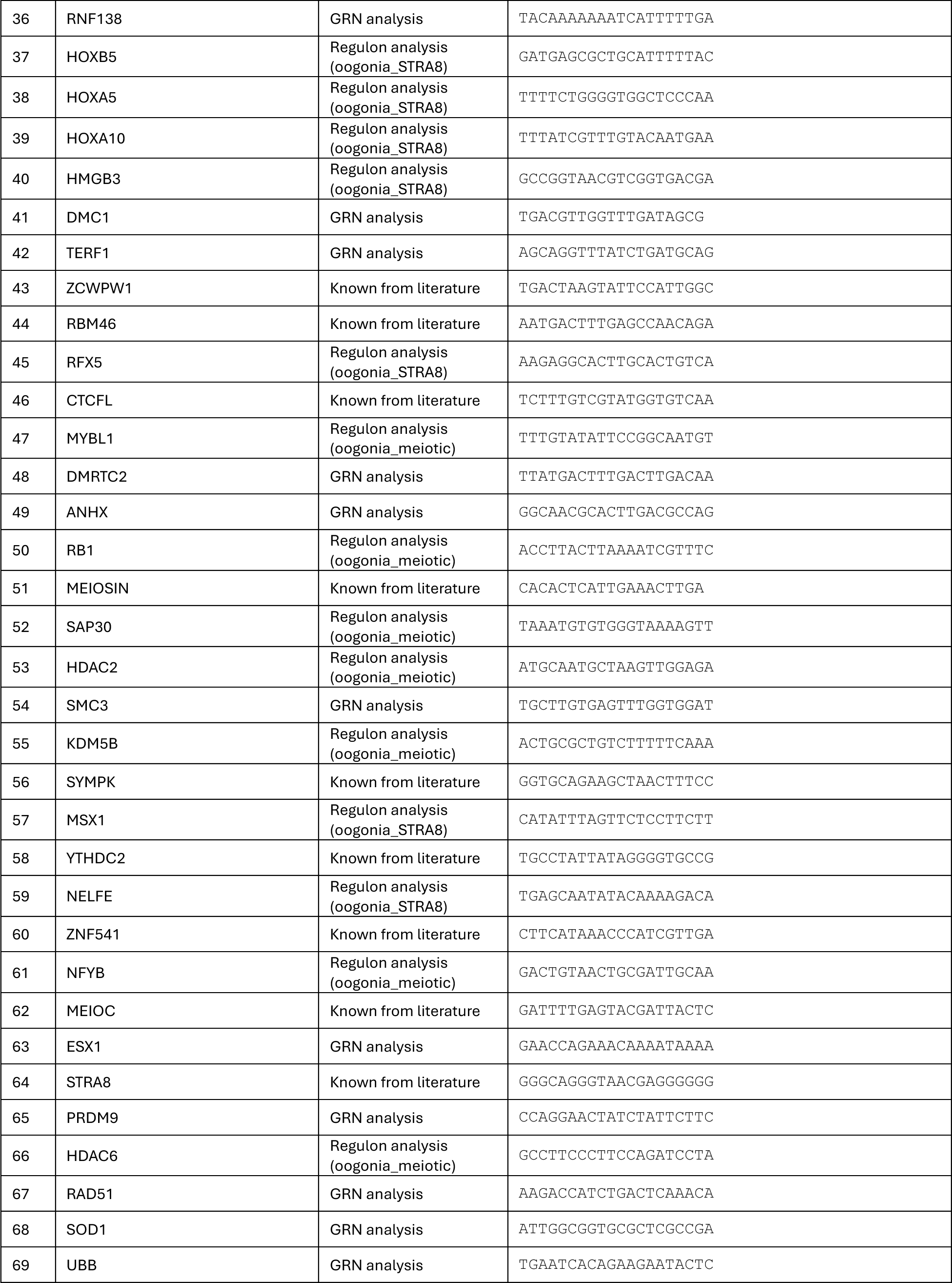
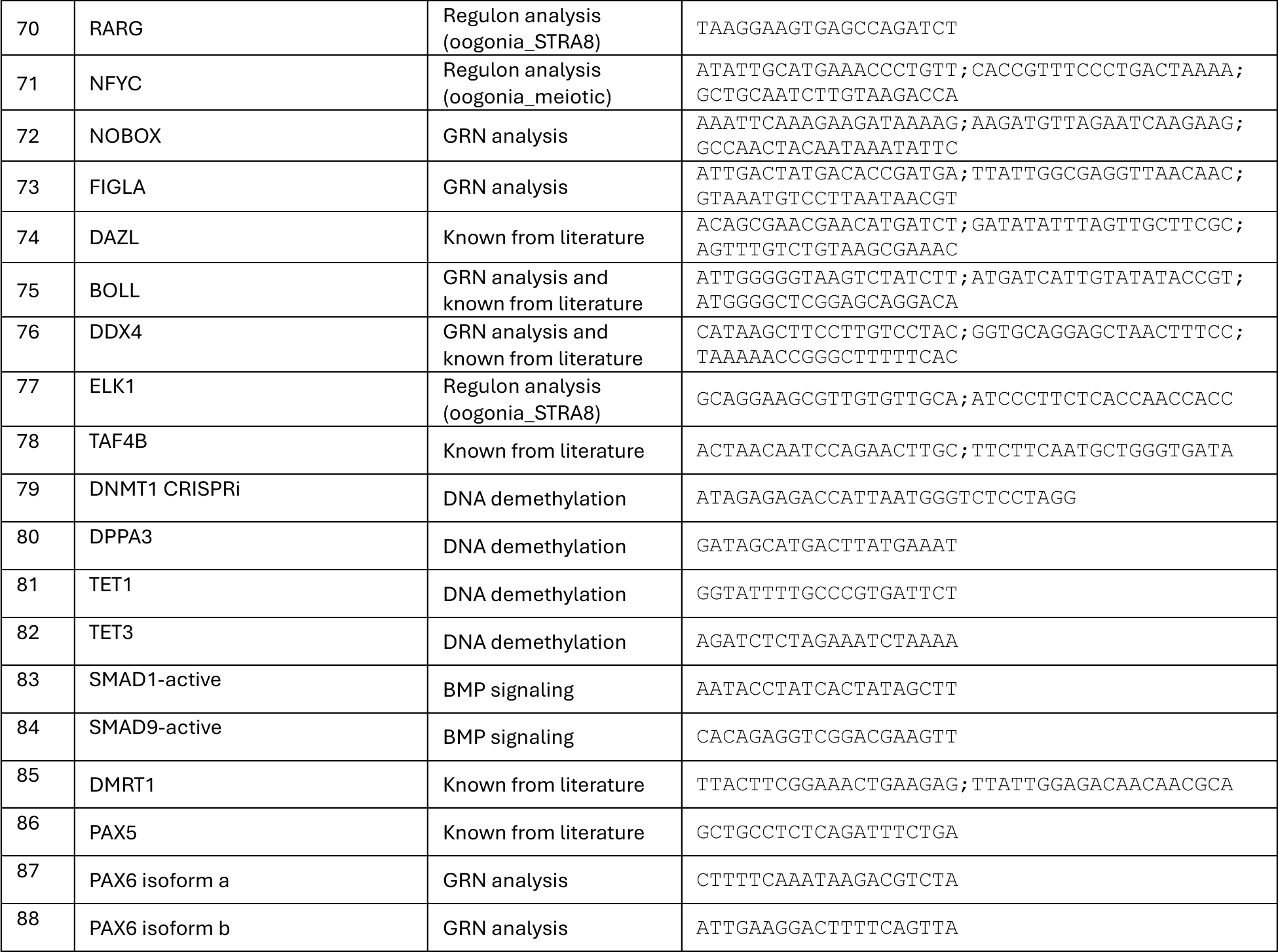
Factors screened for meiosis induction. #1-78 were included in barcode enrichment screening, and #79-88 were added for scRNAseq screening.

**Supplementary Table 3.**
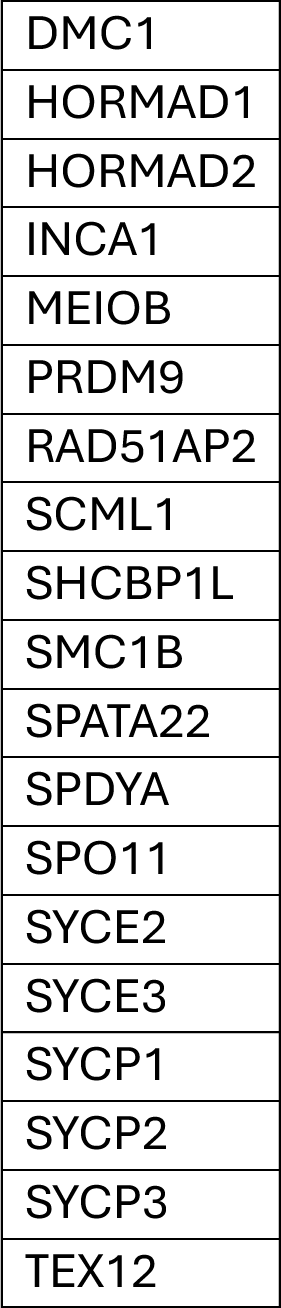
Genes used for meiosis gene score calculation.

**Supplementary Table 4.**
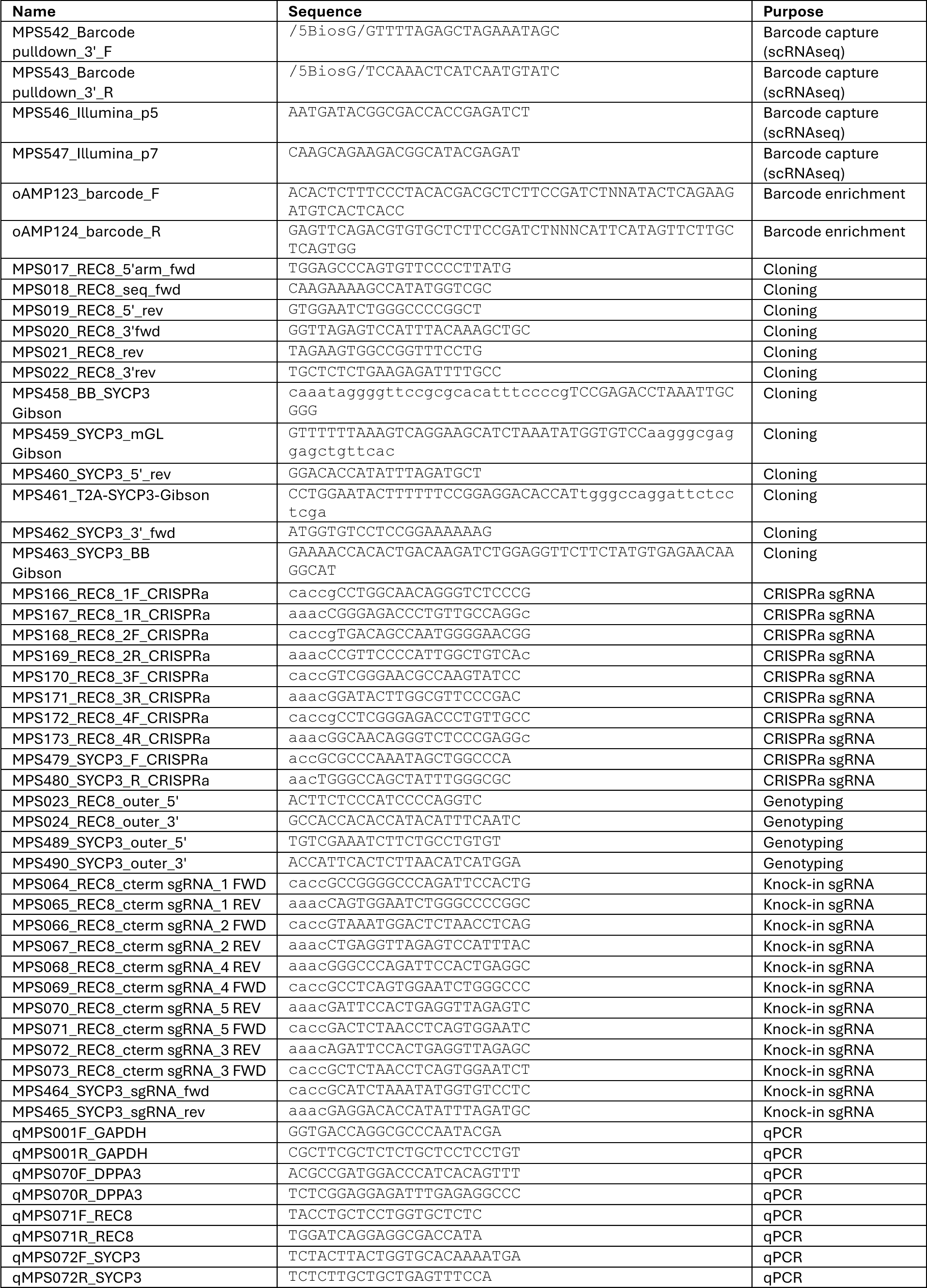
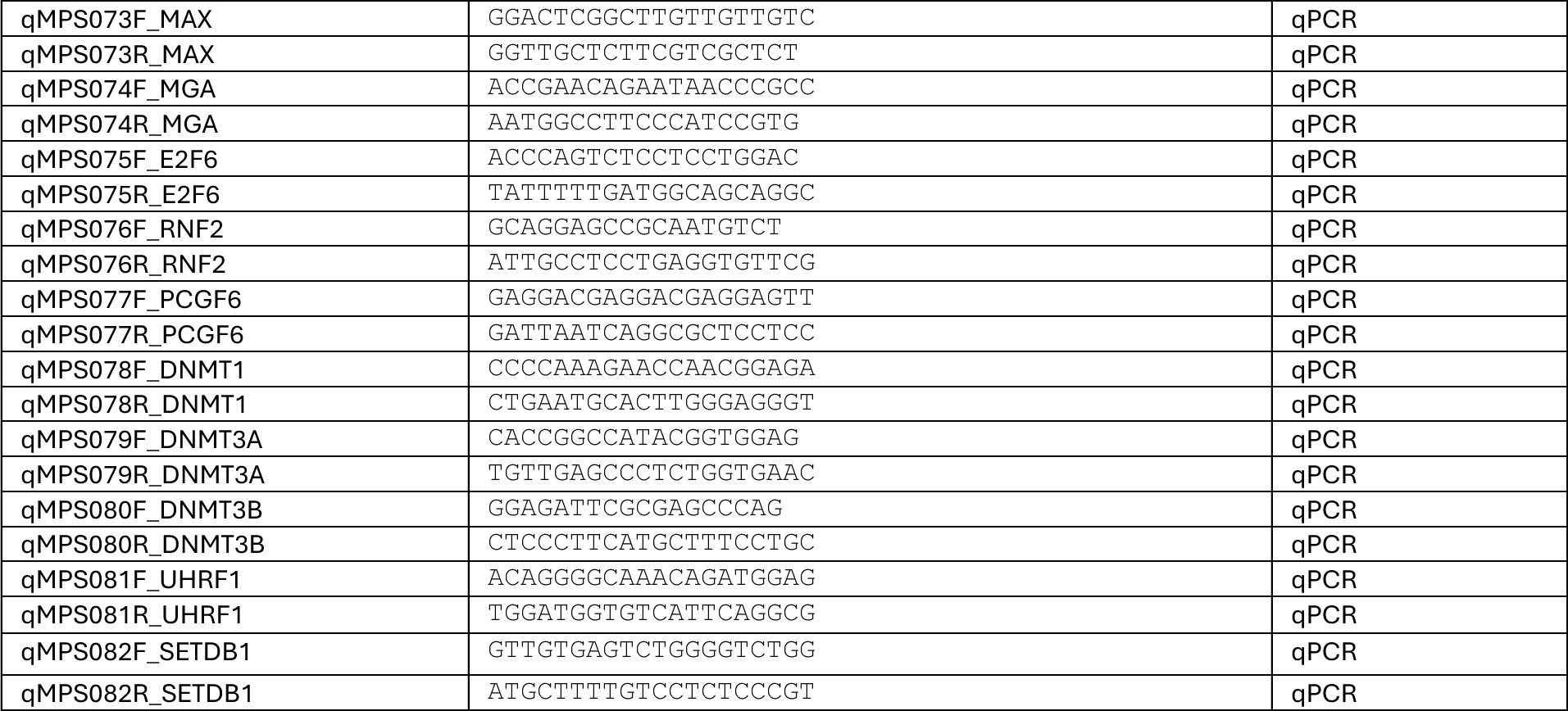
Oligos used in this study.

**Supplementary Table 5.** Antibodies used in this study.

| Primary Antibodies |  |  |  |  |  |
| --- | --- | --- | --- | --- | --- |
| Target | Antibody type | Supplier | Catalog# | Dilution | RRID |
| SYCP3 | Rabbit IgG, polyclonal | Abcam | ab15093 | 1:250 | AB_301639 |
| DDX4 | Rabbit IgG, polyclonal | Abcam | ab13840 | 1:250 | AB_443012 |
| KI67 | Rat IgG, monoclonal | Thermo | 14-5698-37 | 1:250 | AB_2865119 |
| HORMAD1 | Rabbit IgG, polyclonal | Proteintech | 13917-1-AP | 1:250 | AB_2120844 |
| γH2AX | Mouse IgG, monoclonal | Millipore Sigma | 05-636 | 1:500 | AB_309864 |
| Centromere | Human IgG, polyclonal, FITC-labeled | Antibodies Inc | 15-235-F | 1:200 | AB_2797147 |
| SYCP3 | Goat IgG, polyclonal | R&D Systems | AF3750 | 1:250 | AB_2197194 |
| T2A | Rat IgG, monoclonal | Millipore Sigma | MABE1923 | 1:250 | AB_3097817 |
| TEX12 | Rabbit IgG, polyclonal | Abcam | ab122455 | 1:250 | AB_11128111 |
| RAD51 | Mouse IgG, monoclonal | Novus | NB100-148 | 1:250 | AB_10002131 |
| Secondary antibodies (all donkey polyclonal) |  |  |  |  |  |
| Target | Fluorophore | Supplier | Supplier Number | Dilution | RRID |
| Goat IgG | AF568 | Thermo Fisher | A11057 | 1:500 | AB_2534104 |
| Mouse IgG | AF647 | Thermo Fisher | A31571 | 1:500 | AB_162542 |
| Rabbit IgG | AF488 | Jackson | 711-545-152 | 1:500 | AB_2313584 |
| Rat IgG | CF568 | Sigma | SAB4600077 | 1:500 | AB_2827516 |
| Rabbit IgG | AF568 | Thermo Fisher | A10042 | 1:500 | AB_2534017 |
| Rat IgG | Dylight755 | Thermo Fisher | SA5-10031 | 3:500 | AB_2556611 |

**Supplementary Table 6.** List of samples in the scRNAseq screening experiment, and numbers of annotated cell types.

| # | Cell line | Factors | Sorting | Total Cells | Germ cell | Germ cell mitotic | PGC | Oogonia STRA8 | Oogonia meiotic | Pre-oocyte | Pre-spermatogonia |
| --- | --- | --- | --- | --- | --- | --- | --- | --- | --- | --- | --- |
| 1 | PGP1 D4TR8G | BCL2, HOXB5, and STRA8 | unsorted | 3553 | 1323 | 350 | 257 | 1605 | 0 | 0 | 18 |
| 2 | F3 D4TS3G | BCL2, HOXB5, and STRA8 | unsorted | 5308 | 1187 | 320 | 299 | 3464 | 4 | 0 | 34 |
| 3 | PGP1 D4TR8G | BCL2, HOXB5, STRA8, and the pool of 88 factors | unsorted | 90809 | 32279 | 10728 | 9246 | 37839 | 27 | 1 | 689 |
| 4 | PGP1 D4TR8G | BCL2, HOXB5, STRA8, and the pool of 88 factors | sorted DDX4+ | 59668 | 20629 | 6630 | 6399 | 24732 | 65 | 6 | 1207 |
| 5 | PGP1 D4TR8G | BCL2, HOXB5, STRA8, and the pool of 88 factors | sorted REC8+ | 76522 | 26398 | 11405 | 9701 | 28595 | 14 | 0 | 409 |
| 6 | F3 D4TS3G | BCL2, HOXB5, STRA8, and the pool of 88 factors | unsorted | 50837 | 14834 | 4481 | 3267 | 27916 | 38 | 1 | 300 |
| 7 | F3 D4TS3G | BCL2, HOXB5, STRA8, and the pool of 88 factors | sorted DDX4+ | 59742 | 14949 | 6030 | 5233 | 32593 | 100 | 4 | 833 |
| 8 | F3 D4TS3G | BCL2, HOXB5, STRA8, and the pool of 88 factors | sorted SYCP3+ | 88342 | 23339 | 6991 | 5549 | 51541 | 238 | 6 | 678 |
| 9 | PGP1 D4TR8G | Pool of 88 factors | unsorted | 10069 | 3607 | 2197 | 2213 | 2019 | 1 | 0 | 32 |
| 10 | PGP1 D4TR8G | Pool of 88 factors | sorted DDX4+ | 28038 | 9798 | 6119 | 5401 | 6617 | 2 | 0 | 101 |
| 11 | PGP1 D4TR8G | Pool of 88 factors | sorted REC8+ | 27982 | 10776 | 5316 | 5497 | 6114 | 6 | 1 | 272 |
| 12 | F3 D4TS3G | Pool of 88 factors | unsorted | 35747 | 9359 | 9648 | 9883 | 6772 | 1 | 0 | 84 |
| 13 | F3 D4TS3G | Pool of 88 factors | sorted DDX4+ | 25380 | 10364 | 4287 | 4396 | 5996 | 11 | 1 | 325 |
| 14 | F3 D4TS3G | Pool of 88 factors | sorted SYCP3+ | 32779 | 12425 | 5816 | 5202 | 8966 | 34 | 0 | 336 |
| 15 | Mix of PGP1 D4TR8G and F3 D4TS3G | No factors | unsorted | 5312 | 1227 | 1880 | 2016 | 189 | 0 | 0 | 0 |

## Notes

### Competing Interest Statement

MPS and GMC have filed a provisional patent application on the meiosis induction protocol. A full list of GMC conflicts of interest can be found at https://arep.med.harvard.edu/gmc/tech.html Other authors declare that they have no competing interests.

